# A division of labor controls the degradation of fucoidans in the ocean

**DOI:** 10.1101/2025.08.13.670036

**Authors:** Andreas Sichert, Shaul Pollak, Akshit Goyal, Otto X. Cordero, Uwe Sauer

## Abstract

Fucoidans—complex polysaccharides produced by brown algae and diatoms—contribute to long-term carbon sequestration due to their resistance to microbial degradation^1,2^. While individual microbes can break down portions of these polymers^3–5^, it remains unclear whether complete breakdown is possible in nature and, if so, by what mechanisms. Here we show that fucoidans are degraded through synergistic interactions between specialized bacteria with conserved metabolic functions. Using metabolomic analysis of a reconstructed marine consortium, we uncovered functional guilds of bacteria that target either the sulfated fucose backbone or the side-branches of rare monomers. This division of labor leads to an unexpectedly high number of positive interactions between different degraders that enhanced degradation efficiency up to 97.1%. Despite variation in fucoidan structure across different types of algae, the metabolic functions of degraders remained conserved, enabling quantitative prediction of degradation outcomes based on community composition. Our findings suggest that the environmental turnover of complex macromolecules depends not only on individual metabolic capabilities but also on ecological interactions shaped by substrate architecture. This work provides a mechanistic framework for understanding carbon cycling in the ocean and for engineering synthetic microbial consortia to degrade recalcitrant polysaccharides.

## Main

Plant and algal polysaccharides are among the most abundant and diverse macromolecules on Earth^6,7^. Their degradation drives carbon cycling^8,9^, promotes gut health^10^, and enables sustainable biotechnologies^11,12^. As the main component of protective extracellular matrices in plants and algae, the chemical diversity of polysaccharides has escalated a co-evolutionary arms race, driving the diversification of carbohydrate-active enzymes and their reshuffling among microbial degraders via horizontal gene transfer^13–16^. While these distributed metabolic capabilities suggest that microbial interactions may significantly influence degradation efficiency, it remains unclear how multiple degraders coexist and engage in synergistic interactions rather than competition^17–22^. Consequently, we lack a quantitative, mechanistic framework linking the metabolism and interactions of individual degraders to degradation on a community level. This gap hinders our understanding of microbial contributions in carbon cycling and our ability to design microbial consortia for efficient degradation of diverse substrates.

In marine ecosystems, brown algae and diatoms produce the heterogeneous polysaccharide fucoidan, making them key players for carbon sequestration. They account for one-fifth of marine primary production and, through sinking biomass and particles, export 5 Gt C yr⁻¹ to the ocean depths, where the carbon can be stored for millennia^23–25^. These natural processes are increasingly harnessed in brown-algal aquaculture, which is projected to contribute at least 0.5% of the 1 Gt CO₂ yr⁻¹ sequestration target set for nature-based climate solutions by 2050^26^. Fucoidan is a major agent of algal carbon export, as it constitutes 25–50% of the cell wall and mucilage^27,28^, promotes particle formation, and resists microbial degradation for up to several months^1,29^. This stability is likely attributable to its complex structure, commonly featuring a backbone of sulfated fucose with varying monomer composition, sulfation, and branching patterns across algal species^27,30^. Because of this complexity, only a few bacterial species are known to degrade fucoidans, and those that do typically encode dozens of fucoidan-active enzymes, yet access only specific structural variants and achieve incomplete degradation^3–5^. Metagenomic studies reveal co-occurring degraders with complementary enzyme repertoires that potentially target different regions of the polysaccharide^31–33^, suggesting that complete degradation may depend on cooperation among specialized microbes—an ecological hurdle that could stabilize fucoidan in a dilute environment like the ocean and help why so many algae rely on this polysaccharide as a protective layer.

### Genomic specialization structures a fucoidan-degrading microbial consortium

To understand the role of microbial interactions in fucoidan degradation, we enriched a bacterial community of from coastal seawater using fucoidan from the common brown alga *Fucus vesiculosus* as the sole carbon source (Supplementary Table 1). In this fucoidan, fucose has an average of 1.8 sulfate groups and accounts for 80% of all monomers, distributed as 50% in the alternating α-1,3/α-1,4-linked main chain and 30% in short side branches (Fig. 1a). The remaining 20% comprises rare sugars such as xylose, galactose, mannose, and glucuronic acid, which form distinct side branches^27,30^. After 12 sequential growth–dilution cycles, enrichments achieved 90% substrate degradation (Extended Data Fig. 1a-c; Supplementary Table 2). Metagenomic analysis recovered 79 metagenome-assembled genomes (MAGs), with Verrucomicrobiota – a phylum known for degraders of complex polysaccharides – dominating the community at over 80% relative abundance. Notably, the final points of all enrichments were dominated by *Luteolibacter*, a close relative of known fucoidan degraders that inhabit algal surfaces^33,34^.

**Figure 1:**
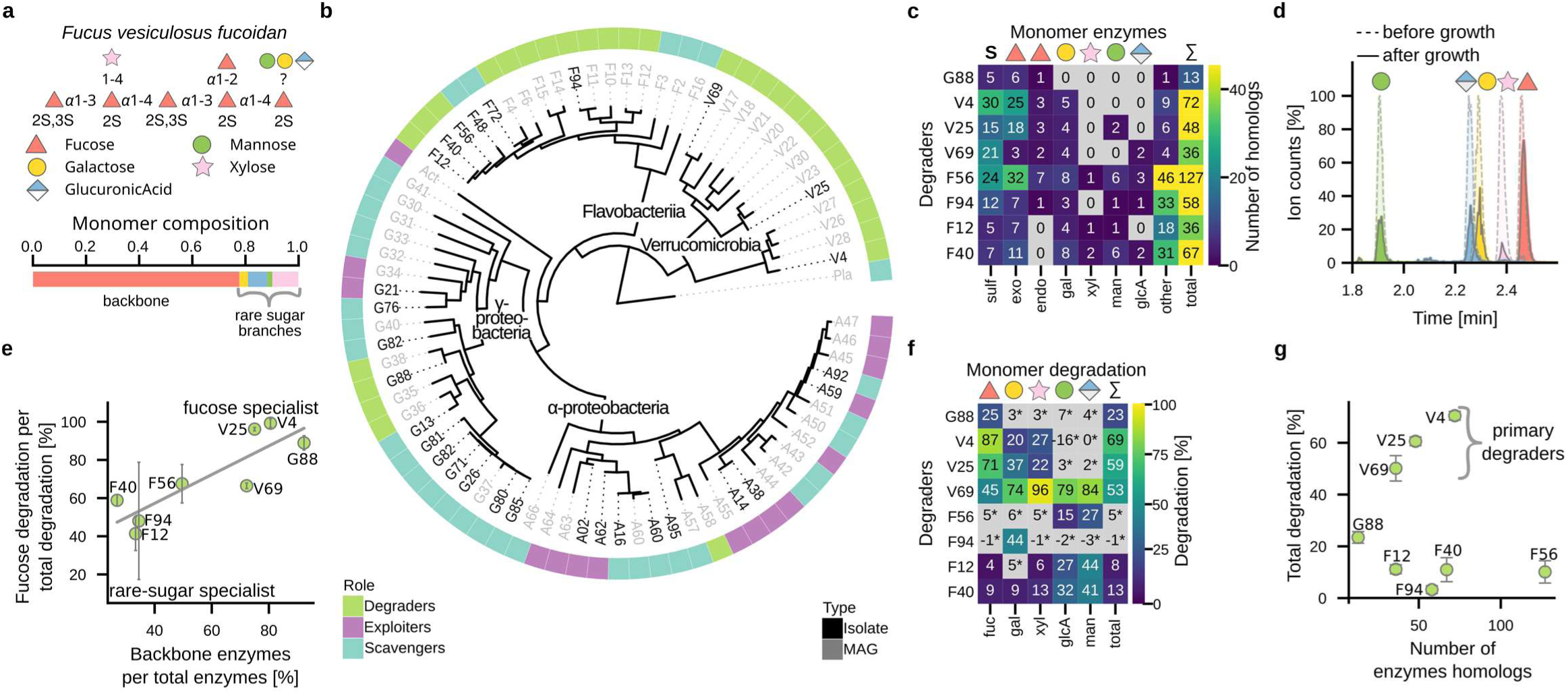
Fucoidan degraders in communities specialize through complementary monomer degradation. **a**, Structure of fucoidan from *Fucus vesiculosus* and corresponding monosaccharide composition shown as bar graph. Monosaccharides are depicted following the Symbol Nomenclature for Glycans; glycosidic linkages are indicated as text labels. **b**, Phylogenetic tree of marine bacterial strains enriched on fucoidan as sole carbon source. The outer ring indicates inferred metabolic roles; names are colored to distinguish isolates from metagenome-assembled genomes. **c**, Number of sulfatases and CAZymes implicated in fucoidan degradation across degraders. Enzyme classes are abbreviated as follows: endo (endo-fucoidanases: GH107, GH168), exo (exo-fucosidases: GH29, GH95, GH141), gal (galactosidases: GH97, GH36, GH117), glcA (glucuronidases: GH115), man (mannosidases: GH92, GH130), and xyl (xylosidases: GH39, GH120). **d**, Representative targeted LC–MS chromatograms of acid hydrolyzed culture supernatants of V69 before (dotted) and after (solid) growth on fucoidan used to infer monomer degradation. Peaks are colored by monomers identified by multiple reaction monitoring and retention time. For visualization, ion counts of monomers are normalized to before growth samples. **e,** Correlation between relative abundance of backbone-degrading enzymes and the proportion of fucose degraded. Points and error bars show mean ± s.d. of three biological replicates. **f**, Heatmap showing the mean monomer degradation obtained from three biological replicates; gray indicates no significant change from control. **g**, Number of fucoidan degrading enzymes versus total fucoidan degradation across degraders. Points and error bars show mean ± s.d. of three biological replicates.

From these enrichments, we established a strain collection that reflects the taxonomic and functional diversity of the community (Fig. 1b). Based on gene content, MAGs were classified into three ecological guilds: i) ‘degraders’, which harbor fucosidases and sulfatases in their genomes and initiate polysaccharide breakdown; ii), ‘exploiters’, which lack these enzymes but possess genes for fucose catabolism, allowing them to compete with degraders for released fucose; and iii) ‘scavengers’, which do not contain either class of enzymes and instead rely on metabolic byproducts for growth (Extended Data Fig. 1d). This classification revealed a surprisingly high diversity of 30 degraders (Fig. 1b). Among them, we isolated V25 (*Luteolibacter*), V69 (*Roseibacillus*), G88 (*Pseudocolwellia*), and four members of the *Flavobacteriia* (F12, F56, F40 and F94). Additionally, we isolated five exploiters and sixteen scavengers, including the previously characterized *Lentimonas* sp. CC4 (V4)^3^. Collectively, these 29 strains provide a foundation for dissecting the mechanisms of fucoidan degradation in microbial communities.

Genome analysis of the isolated degraders revealed an unexpectedly large and diverse repertoire of fucoidan-degrading enzymes. We identified 35 genomic polysaccharide utilization loci (PULs) associated with fucoidan degradation (Extended Data Fig. 2a; Supplementary Table 3). Within these regions, canonical “backbone” enzymes targeting sulfate and fucose^3,4,35^ co-localized with putative “debranching” enzymes belonging to 44 CAZyme families, some of which act on rare sugars. Each degrader encoded 12 to 127 different enzymes, defined using a 60% identity threshold, with an average pairwise overlap of only 9% amounting to 547 unique enzymes across all genomes of the enrichment cultures (Fig. 1c; Extended Data Fig. 2b,c). Strikingly, all enzymes of strain G88 were encoded on a 112 kb plasmid, while strain F56 harbored a 623 kb genomic island containing five PULs (Extended Data Fig. 3). These observations are consistent with the lateral acquisition of large CAZyme-rich gene clusters^13,16^. Together, these findings indicate that enzyme repertories are shaped by gene mobility and partial sampling from a vast environmental pool, culminating in a highly diverse enzymatic landscape for fucoidan degradation.

Despite the unique nature of enzyme repertoires across different genomes, a clear functional distinction emerged, differentiating debranching from backbone-cleaving mechanisms. Independent of the total enzyme count, the proportion of all backbone-targeting enzymes per total fucoidan acting enzymes varied significantly – ranging from under 30% in F40 to over 90% in G88 (Extended Data Fig. 4) – indicating varying levels of genomic specialization for monomer degradation. The fucose specialists V25, V4, and G88 lacked glucuronidases and xylosidases, suggesting a limited capacity to cleave rare side-chain sugars (Fig. 1c). Conversely, the rare-sugar specialists F12, F40, and F94 lacked endo-acting fucoidanases, which are required cleave the main chain. Generalist strains, such as V69 and F56, encoded a broader array of enzymes, encompassing both functional roles. These specialized enzymatic repertoires indicate distinct monomer preferences among the degraders.

### Monomer degradation reveals functional complementarity among degraders

We validated our genomic predictions about enzymatic specialization using a targeted liquid chromatography–mass spectrometry (LC-MS) assay to measure degradation at the monomer level. This degradation was inferred by comparing the concentrations of monomers bound within fucoidan before and after growth (Fig. 1d; Eq. 1,2), released through acid hydrolysis of culture supernatants. Monomers were quantified using established derivatization protocols and an optimized 3.5-minute LC-MS method^36,37^. As in previous studies^3,4^, this approach effectively captured the combined effect of enzymatic cleavage of the polysaccharide and microbial uptake of the released monomers—a process we refer to as monomer degradation— which could be quantified across a range from 5% to 99.9% (Extended Data Fig. 5; Supplementary Table 4). To confirm metabolic specialization, we evaluated each degrader’s monomer preference by determining the fraction of degraded carbon derived from fucose, revealing a strong correlation with the genomic proportion of backbone-cleaving enzymes in the genome (*r* = 0.75, *P* < 0.001; Fig. 1e). These results confirm the predicted specialization for fucose, or rare monomers based on genomic analysis and underscores that monomer metabolism is a key feature that differentiates fucoidan degraders.

Unexpectedly, however, none of the degraders were able to completely break down any specific monomer type, even those aligned with their apparent specialization (Fig. 1f). The fucose specialists V4 and V25 achieved only 87% and 71% fucose degradation and low amounts of galactose and xylose, but were unable to degrade mannose and glucuronic acid. These monomers were solely degraded by F12, F40, F56, and V69, which reached up to 86% rare monomer degradation, but showed markedly lower fucose degradation than the dedicated fucose specialists. These limited metabolic capacities resulted in total degradation of only 69%, 59%, and 53% for the strong primary degraders V4, V25, and V69, respectively, which is substantially lower than the 90% achieved in enrichment cultures. Notably, the total degradation efficiency among isolates did not correlate with the total number of enzymes (Fig. 1g), highlighting the difficulty of inferring total degradation from genomes. For instance, although F56 encodes over a hundred enzymes, it achieved almost negligible total degradation, whereas the metabolically specialized G88 achieves 25% fucose degradation with a mere 12 enzymes. Together, these findings suggest that complete fucoidan degradation may require a division of labor among bacteria with complementary metabolic capabilities.

### Synergism in pairwise co-cultures

To assess the impact of microbial interactions on fucoidan degradation, we measured the fold changes in biomass and total degradation for the primary degraders V4, V25, and V69 when co-cultured individually with each of the 29 strains, comparing the results to monoculture controls (Fig. 2a; Extended Data Fig. 6; Supplementary Table 5). Of the 87 co-culture pairs tested, 33 exhibited significant changes which broadly aligned with predictions based on genome-based guild classifications: Eight pairs with scavengers increased biomass without influencing degradation, supporting their role in recycling byproducts that are not directly linked to the degradation pathway. Conversely, seven pairs with predicted exploiters reduced both biomass and fucoidan degradation, suggesting competition for common degradation products, particularly fucose. Remarkably, 16 degrader–degrader co-cultures showed positive interactions, characterized by increases in both biomass and degradation, as well as higher growth rates. The only exception was G88, which increased degradation when paired with V69, but acted as an exploiter when paired with V4 and V25. These results show that rather than competition, degraders engage in cooperative interaction that are crucial for achieving complete fucoidan degradation.

**Figure 2:**
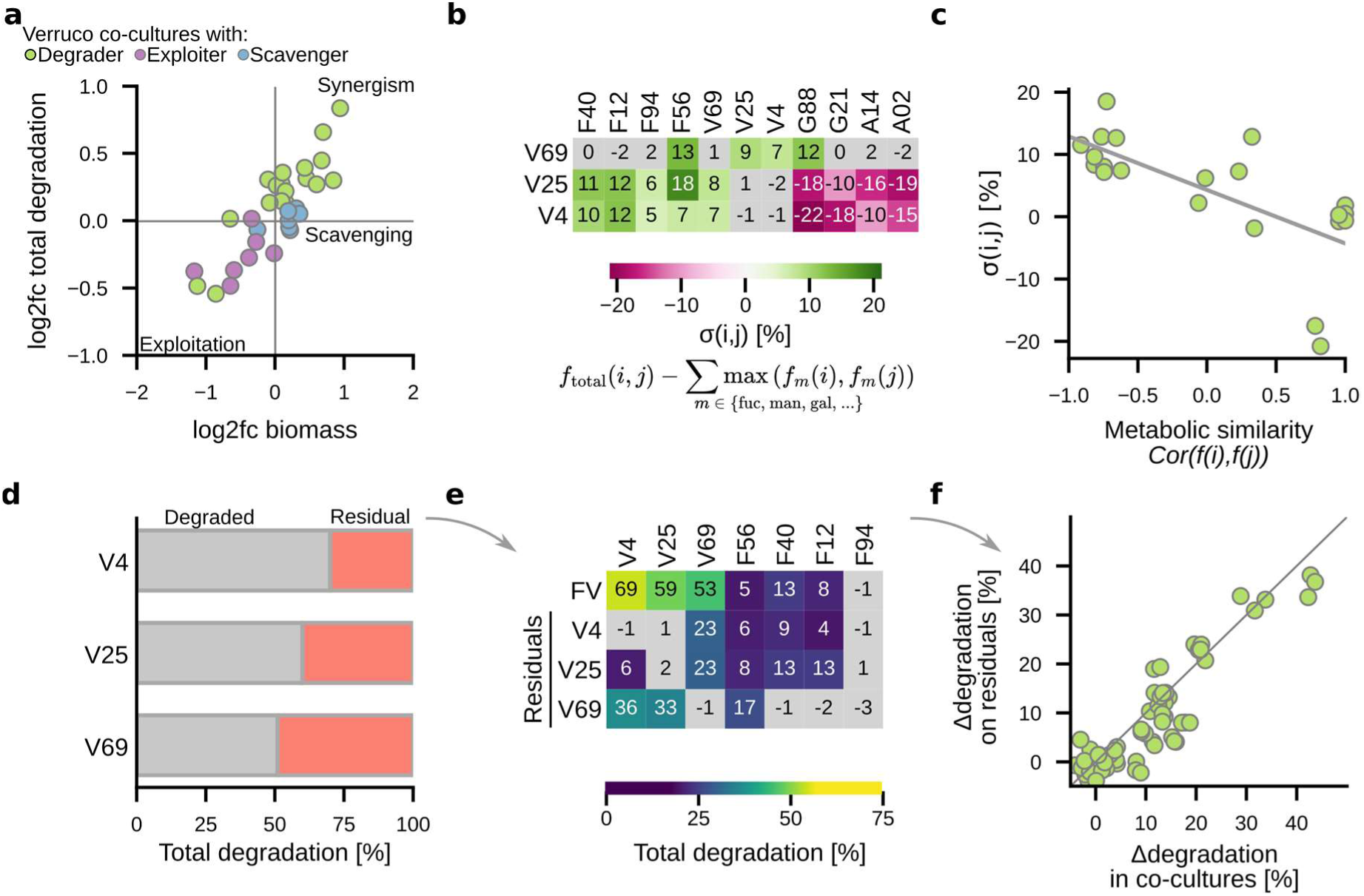
Synergistic interactions between complementary degraders in pairwise co-cultures. **a**, Fold-changes in biomass and total fucoidan degradation in 87 pairwise co-cultures of Verrucomicrobiota strains with other community members, relative to Verrucomicrobiota monocultures. Each point is color-coded by the metabolic role of the partner strain (degrader, exploiter, or scavenger). **b**, Heatmap showing the synergism score between Verrucomicrobiota strains and other community members. Synergism score is calculated from a null expectation assuming no interspecies interaction. **c**, Correlation between synergism score and metabolic similarity of strain pairs, calculated from the correlation of normalized monomer degradation profiles from monocultures. **d**, Bar graph showing the fraction of fucoidan degraded by three primary degraders, and the residual substrate purified for cross-feeding assays. **e**, Heatmap showing the effective degradation of residual fucoidans by secondary degraders. Effective degradation was calculated as the product of the residual substrate fraction left by the primary degrader and the degradation of the corresponding purified residual fucoidan by the secondary degrader in monoculture. Data in panels **a–e** represent the mean of three biological replicates. **f**, Correlation between the additional degradation observed in co-cultures (relative to the respective primary degrader alone) and the effective degradation achieved by secondary degraders on residual fucoidan. All individual data points from three biological replicates are shown.

The observed positive interactions among degraders could not be explained solely by resource partitioning, indicating the emergence of a synergistic division of labor. We quantified a synergism score, denoted as *σ(i,j)*, by calculating the difference between observed total degradation and a null hypothesis that assumed no interaction beyond mere resource partitioning (Fig. 2b; Eq. 3). The strongest synergism of 18% occurred between V25 and F56, which together achieved 79% total degradation. The highest degradation levels overall were found in the pairs V25/V69 (82%) and V4/V69 (93%), each exhibiting synergistic contributions of about 8%. Overall, a strong negative correlation was discovered between the calculated synergism *σ(i,j)* and the dissimilarity of monomer degradation profiles between paired strains (*r* = – 0.79, *P* < 0.001; Fig. 2c), indicating that complementary metabolic capabilities amplified degradation in a non-additive manner. Additionally, we identified positive association between *σ(i,j)* and the difference in the genomic proportion of backbone enzymes relative to total enzymes (r = 0.53, *P* = 0.02; Extended Data Fig. 7). Collectively, these results indicate that synergism emerges when enzymatic and metabolic capabilities are maximized through a division of labor between fucose and rare monomer specialists.

To determine whether the observed synergistic resulted from resource-mediated interactions or other metabolic exchanges among degraders, we investigated cross-feeding using purified residual fucoidan (>1 kDa) obtained from stationary-phase Verrucomicrobiota cultures (Fig. 2d). Using these residuals as the sole carbon source, we assessed their degradation by seven ‘secondary’ degraders using LC-MS (Fig. 2e). All secondary degraders, except F94, were able to utilize at least one residual substrate. The additional degradation of residuals by these secondary degraders quantitatively aligned with the enhanced performance observed in their corresponding co-cultures (R² = 0.90, *P* = < 0.001; Fig. 2f; Eq. 4), demonstrating that synergism can occur independently of direct cell–cell contact or the exchange of signalling molecules. Furthermore, we observed that F56 has a threefold increase in residual degradation compared to its performance in monoculture (Fig. 2e). This suggests that the prior degradation of certain moieties likely facilitated F56’s enzymatic activity by removing moieties that would otherwise impede its function. Conversely, strains F12 and F40 exhibited unchanged degradation capabilities, suggesting that their roles are fully complementary to those of V25 or V4. In contrast, the primary degraders showed a decrease in performance, likely due to overlapping substrate specificities.

### Quantitative prediction of synergistic degradation from metabolic data

To explore whether cooperative effects extend beyond pairs, we measured degradation across all 127 combinations of the seven consistent degraders, excluding G88. Total degradation ranged from 4% to 97.1% and generally increased with community richness (Fig. 3a). One striking example of emergent cooperation was observed between strains F56 and F94, which demonstrated minimal degradation in monoculture but achieved 50% degradation when combined (Extended Data Fig. 8a). Overall, the marginal contribution of each degrader declined as the overall performance of the background community increased, aligning with the concept of growing functional redundancy when saturation is approached. Remarkably, even under these saturated conditions, 80% of interactions were still synergistic beyond what would be expected from simple resource partitioning, while negative interactions were infrequent and weak (Eq. 3; Fig. 3b). These findings suggest that most interactions among degraders were either additive or cooperative, indicating a smooth structure-function landscape with minimal higher-order effects and an unexpected degree of predictability in community function^38^.

**Figure 3:**
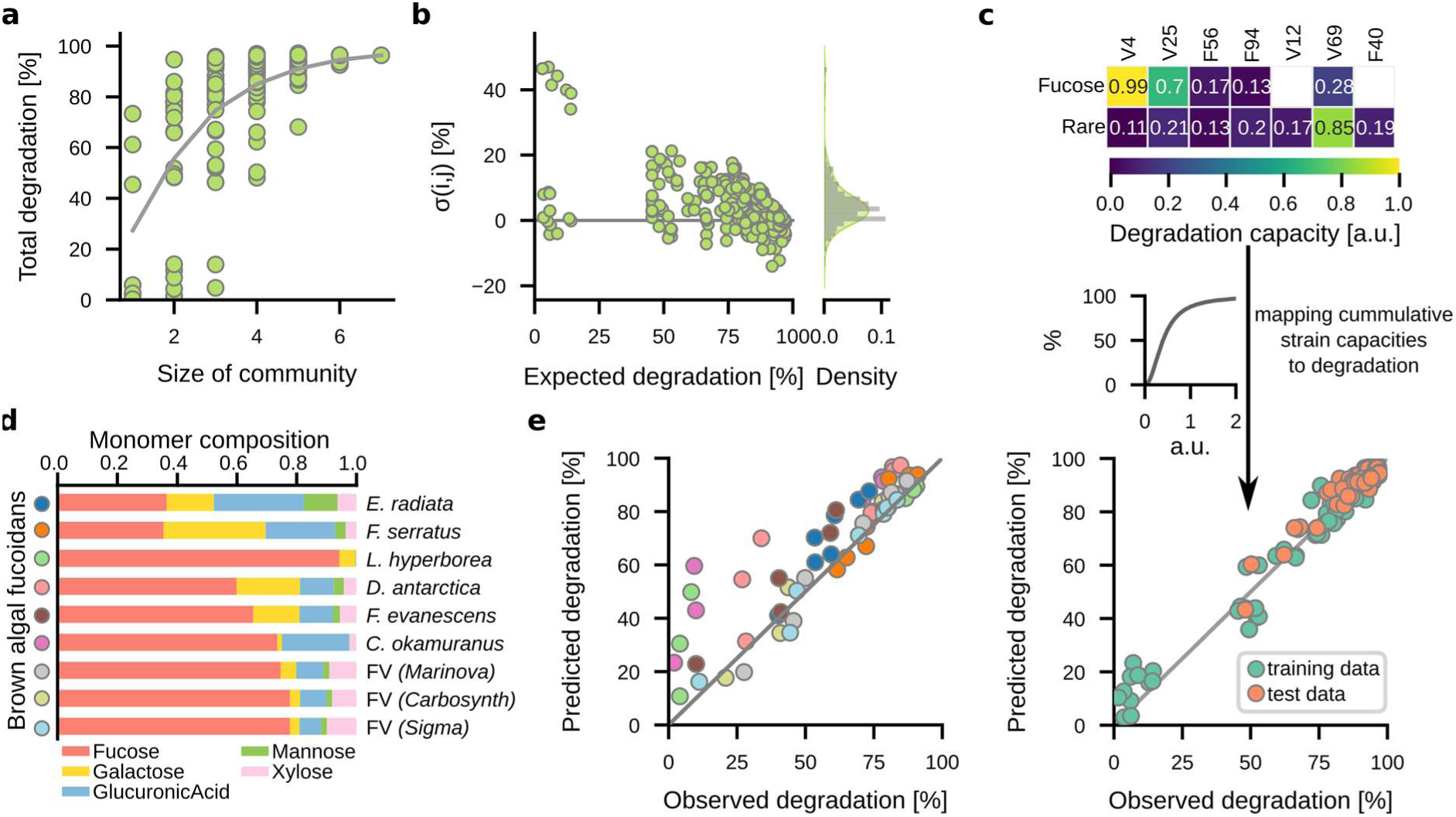
Division of labor enables predictable degradation synergies across different fucoidan structures. **a**, Total fucoidan degradation in all 127 possible combinations of up to seven degraders. Data points show the mean of three biological replicates; the grey line denotes the mean degradation per community size. **b,** Synergism score in 127 communities compared to expected degradation under a null model assuming no interactions. Left: scatter plot of expected degradation versus synergism score. Right: density plot of synergism score distribution. All individual data points from three biological replicates are shown. **c**, Predictive model of community degradation based on strain-specific substrate preferences. Top: heatmap of inferred degradation capacities for fucose and rare sugars across seven strains. Center: non-linear Hill function mapping cumulative strain capacity to predicted degradation. Bottom: predicted versus observed degradation across 127 communities; points are colored by training or test set and show the mean of three biological replicates. **d**, Monosaccharide composition of nine distinct brown algal fucoidans. **e**, Observed versus predicted degradation of nine fucoidan substrates across seven selected communities. Points are colored by substrate identity and show the mean of three biological replicates.

To further assess the predictability of degradation in complex communities, we developed a mechanistic model that links community composition to degradation output (Fig. 3c). This model was inspired by the observed resource partitioning between fucose and rare monomer degraders and represents fucoidan as two monomer pools, where strains are characterized by their ability to degrade each type (Eq. 5). Given that degradation tended to saturate in richer communities, we hypothesized that community-level degradation could be predicted by a nonlinear combination of the capabilities of individual strains, captured through a Hill function (Eqs. 6 and 7). Model parameters—strain-specific degradation potentials and Hill coefficients—were inferred from data on one-, two-, and three-member communities (Extended Data Fig. 9). This straightforward framework proved to be highly effective, accurately predicting degradation across all 127 community combinations (R² = 0.96, *P* < 0.001; Fig. 3c; Supplementary Table 6), including the emergent interaction between F56 and other degraders. These results highlight that, despite the structural complexity of fucoidan and the diversity of enzymatic pathways involved, community-level degradation can be reliably predicted from the synergistic contributions of individual strains to fucose and rare monomer utilization.

The nearly complete absence of higher-order interactions and the functional redundancy among degraders suggest that maximal degradation can be achieved through various community configurations. Some configurations depended on a few potent strains (e.g., V69 paired with V25), while others functioned based on synergistic combinations of individually weaker degraders (Extended Data Fig. 8b). For example, four *Flavobacteriia* strains, which, although ineffective alone, collectively approximated the degradation performance of V69 by cooperatively targeting both fucose and rare monomers. This functional redundancy implies that diverse communities can serve as a buffer against the stochastic fluctuations that often occur in species composition typical of particle-associated marine microbiomes^39,40^. However, it also highlights the potential for what we term “diversity-limited” degradation. In such cases, microbial assemblages may lack the complementary degraders necessary to fully exploit available polysaccharide substrates, resulting in suboptimal degradation performance.

Given that the monomer composition of fucoidan varies substantially between brown algal species and across seasons^41^, we investigated whether the principles we discovered above extend to structurally divergent substrates. To this end, we tested seven different communities on nine different types of fucoidans where fucose ranged 35% in *Ecklonia radiata* to 94% in *Laminaria hyperborea* (Fig. 3d), requiring distinct sets of specialized enzymatic activities^3,27^.

The differences in monomer composition strongly influenced the degradation performance of strains in line with their specialized enzymatic and metabolic capabilities. Most notably, strains F12, F40, and F94, which encode a diverse repertoire of debranching enzymes, underperform on the main *Fucus vesiculosus* fucoidan, yet achieve high growth and degradation substrates enriched with rare monomers (Extended Data Fig. 10). Remarkably, without any refitting, our predictive model generalized well to these chemically distinct fucoidans (R² = 0.87; *P* < 0.001; Fig. 3e). Using the parameters trained on fucoidan from *F. vesiculosus,* accurately predicted degradation across all nine substrates by accounting solely for the stoichiometry of fucose and rare monomers. These out-of-sample predictions demonstrate an extraordinary degree of accuracy, considering the complexity of the degradation pathways involved. This suggests that the metabolic capabilities of the degraders and their synergistic interactions remain conserved across different substrates, despite variation in overall monomer composition and polysaccharide structure.

## Discussion

Our results demonstrate that positive interactions among complementary degraders are both frequent and essential for the complete breakdown of fucoidan by marine microbial communities. This emphasizes that its degradation is fundamentally a collective effort. Despite being a particularly complex and variable class of polysaccharides - with dozens of glycosidic linkages and an expansive enzymatic space - the degradation of fucoidans at the community level proves remarkably predictable. This paradox is resolved by a division of labor among degraders, in which individual strains consistently specialize in either the fucose-rich backbone or the rare-monomer side chains, exhibiting conserved metabolic roles across diverse fucoidan structures. This allows for a quantitative mapping of community composition to degradation outcomes and underscores that the co-evolutionary arms race between the protective extracellular matrix of brown algae and microbial degraders unfolds at the monomer level.

A central open question remains regarding the evolutionary and ecological constraints that foster this division of labor. Several non-mutually exclusive hypotheses merit consideration. One possibility involves a metabolic trade-off between fucose and other hexoses: whereas side-chain sugars typically feed into upper glycolysis, fucose enters metabolism at the level of lower glycolysis, requiring gluconeogenic flux to generate upstream intermediates. This mismatch may favour the evolution of specialists that target either the fucose backbone or the side chains, avoiding potentially wasteful cycling in central carbon metabolism^42,43^. Another contributing factor may be the metabolic cost of maintaining and regulating a broad enzymatic arsenal. Fully self-sufficient degraders must coordinate the expression of hundreds of genes, many of which are needed to process low-abundance monomers, making this strategy energetically inefficient. Lastly, the continual loss of genes through neutral processes such as genetic drift, combined with the ability of coexisting degraders to compensate for each other’s deficits, suggests that complete degraders may be evolutionarily unstable. In contrast, gene loss may drive the recurrent emergence of complementary types that together achieve full degradation.

Our findings have implications for both microbial ecology and biotechnology. First, they highlight the challenges associated with engineering single strains to degrade chemically complex substrates such as fucoidan, where identifying precise enzyme functions proves difficult. In contrast, harnessing microbial communities with pre-evolved metabolic complementarity presents a powerful and scalable alternative for biomass conversion. Thus, our work establishes a conceptual and practical framework to process increasingly important brown algal biomass and might extent to other polysaccharides with similar structures such as xylans^20^ and mucins^22^. Second, from an ecological perspective, the necessity for coordinated multi-species activity may limit fucoidan degradation in natural environments. The low abundance of fucoidan-degrading taxa^1,3,31^ and narrow metabolic capabilities suggest that successful degradation may hinge on the stochastic co-occurrence of complementary traits. This reliance on diverse partners may contribute to the long residence times and emerging role in carbon sequestration of fucoidan in the ocean^1^. Our research highlights an underrecognized link between microbial diversity, substrate turnover, and global biogeochemical cycles. In light of the ongoing biodiversity loss^44,45^, this relationship may ultimately influence the ocean’s future capacity for carbon sequestration.

## Materials and Methods

### Chemicals and reagents

Fucoidan from *Fucus vesiculosus* was obtained from Sigma-Aldrich (F8190). Sources for other fucoidans were *Fucus vesiculosus* (Marinova FVF2021547), *Fucus vesiculosus* (Biosynth YF57714), *Fucus serratus* (Biosynth YF09360), *Fucus evanescence* (OceanBasis), *Cladosiphon okamuranus* (Biosynth YF146834), *Durvillea potatorum* (Biosynth YF157165), *Ecklonia radiata* (Biosynth YF157166) and *Laminaria hyperborea* (TheFucoidanStore LowEndo Fucoidan). For derivatization, 1-phenyl-3-methyl-5-pyrazolone (PMP) was obtained from Sigma-Aldrich (M70800). Internal standards for LC–MS analysis included D-galactose-¹³C₆ (Sigma-Aldrich, 605379), D-mannose-¹³C₆ (Sigma-Aldrich, 592994), and PMP-[d₅] (CAS 1228765-67-0), custom-synthesized by BOC Sciences (Shirley, NY, USA). LC–MS-grade solvents and reagents were acetonitrile and methanol (Honeywell), ethanol (Sigma-Aldrich), formic acid (Sigma-Aldrich), and ammonium formate (Merck). Ultrapure water was produced using a Q-POD system (Merck). Unless otherwise stated, all other chemicals were of analytical grade and sourced from Sigma-Aldrich.

### Bacterial growth media

Throughout this study, three distinct media detailed in Supplementary Table 1 were used for i) enrichment of bacteria, ii) isolation of bacteria on solid media, and iii) routine cultivation of bacterial isolates in MBL medium^40^. All media mimicked the ionic composition of coastal seawater and contained 340 mM NaCl, 15 mM MgCl_2_, 6.75 mM KCl and 1 mM CaCl_2_. The pH was buffered pH=8.0 by either bicarbonate in both the enrichment and plate media or 50 mM HEPES in MBL medium. Nutrients were ammonium chloride, sodium phosphate, sodium sulfate, trace metal mix and vitamin mix^40^. Enrichment media contained 0.02% (w/v) fucoidan from Fucus vesiculosus. Solid media was prepared with 2% (w/v) carrageenan (Sigma C1013) and a mix of carbon sources (Acetate, Citrate, Xylose, Galactose, Mannose, Glucose, Fucose, Cellobiose, Tryptone) at 0.002% (w/v) each.

### Enrichment of fucoidan degrading communities

Surface seawater was collected on 30 March 2019 from a rocky shoreline covered with the brown algae *Fucus vesiculosus*, *Ascophyllum nodosum*, and *Laminaria saccharina* near the Marine Science Center of Northeastern University (Canoe Beach, Nahant, Massachusetts, USA; 42°25′10.8732″ N, 70°54′25.686″ W). The seawater was pre-filtered through a 10 μm PTFE membrane filter (Millipore, JCWP04700) and diluted 1:100 to inoculate three replicate enrichment cultures (25 mL each) containing 0.02% (w/v) Fucus vesiculosus fucoidan in 150 mL glass bottles sealed with rubber stoppers. Cultures were incubated at 20 °C in the dark and agitated at 50 rpm.

Enrichment cultures were monitored every 24–48 h for microbial growth (OD₆₀₀) and fucoidan degradation, quantified via the phenol–sulfuric acid method. For each measurement, 200 µL of culture was mixed with 1 mL concentrated sulfuric acid and 200 µL of 5% (v/v) phenol, then incubated at 50 °C for 20 minutes. Absorbance was measured at 490 nm, and fucoidan concentration was determined using an external standard curve of fucose. After 10 days, cultures reached 40–60% degradation of the initial fucoidan amount, at which point serial growth-dilution cycles were initiated by transferring 1:50 into freshly prepared medium every two days for a total of 12 cycles. At every second time point, 10 mL of culture was filtered onto a 0.22 μm Sterivex filter (Millipore, SVGPB1010) for DNA extraction using the DNeasy Blood & Tissue Kit (Qiagen) and subsequent metagenomic sequencing. Final enrichment communities were cryopreserved at −80 °C with 15% (v/v) glycerol.

### Isolation of bacterial strains

From each enrichment culture, 100, 1000, and 10000 cells were plated in triplicate on solid medium in 150 mm Petri dishes (VWR 391-0616) and incubated for 14 days at ambient temperature in the dark. A total of 768 colonies were picked and re-streaked at least three times until only a single colony morphotype was observed. Colonies were lysed in 0.1% (v/v) Triton X-100 in TE buffer for Sanger sequencing of the 16S rRNA gene. Pure isolates were grown in MBL medium containing a mixed carbon source and cryopreserved at −80 °C in 15% (v/v) glycerol. To de-replicate repeatedly isolated strains, we selected 96 isolates based on their 16S sequences for genomic DNA extraction using the DNAadvance kit (Beckman Coulter) and draft genome sequencing. To identify redundant isolates, pairwise average nucleotide identity (ANI) was computed using OrthoANIu v1.2^46^ yielding 28 unique bacterial strains.

### Sequencing of metagenomes and isolate genomes

All sequencing work was carried out in collaboration at the BioMicroCenter at MIT. Metagenomes of enrichment cultures were sequenced with an Illumina NovaSeq6000 S4 flow cell with 150 nt paired end reads. Metagenomes were assembled using SPAdes v3.13.0^47^ with the parameters --meta --only-assembler -k 21,33,55,77. Metagenome-assembled genomes (MAGs) were reconstructed using CONCOCT v1.0.0^48^, MaxBin v2.2.7^49^, and DAS Tool v1.1.2^50^ with default settings. Draft genomes of isolates were sequenced on an Illumina NextSeq 500 platform (150 nt paired-end reads, ∼200× coverage) and assembled using SPAdes v3.13.0 with the parameters --only-assembler -k 21,33,55,77 --careful. To generate closed genomes for selected degraders (Flavo12, Flavo40, Flavo56, Flavo94, Gamma88, Verruco25, and Verruco69), genomic DNA was extracted from 1 mL of culture using the DNeasy Blood & Tissue Kit (Qiagen), and long-read sequencing was performed on Nanopore PromethION FLO-PRO002 flow cells. Hybrid assemblies combining Illumina short reads and Nanopore long reads were generated using Unicycler v0.4.8^51^ with the parameters --keep 3 --mode normal --min_fasta_length 1000 --kmers 77, yielding circularized genome assemblies. Completeness and contamination of MAGs, draft genomes, and closed genomes were assessed using CheckM v1.1.2^52^, resulting in a total of 79 high-quality genomes with accession numbers listed in Supplementary Table 2. Taxonomic classification and phylogenetic reconstruction of isolate and metagenome-assembled genomes were performed using GTDB-Tk v2.0.0^53^ (release 202).

### Annotation of bacterial genomes

Open Reading Frames were predicted and annotated with DRAM v1.2.4^54^. Carbohydrate active enzymes (CAZymes) were identified by HMMER^55^ version 3.3 with hidden Markov models from the dbCAN database^56^ (dbCAN-HMMdb-V10). HMM hits were validated via diamond v0.9.14.115 blastp (--more-sensitive) against the CAZy database (accessed September 2022), retaining only matches with bit scores >100. Sulfatases were identified by hmmsearch against PF00884 (sulfatase domain), using an e-value threshold of <1e−4 and a minimum alignment length of 100 amino acids. Sulfatases were further classified into subfamilies based on diamond blastp (--more-sensitive) against the Sulfatlas v1.2. database^57^.

### Classification of genomes into metabolic roles

We classified genomes into three metabolic roles, degraders, exploiters, and scavengers, based on the presence or absence of key enzymes involved in fucoidan and L-fucose metabolism. Degraders were defined as genomes encoding at least five enzymes from a curated set of ten families of fucoidanase families^3^, comprising glycoside hydrolases (GH29, GH95, GH141, GH107, GH168) and sulfatases (S1_15, S1_16, S1_17, S1_22, S1_25). Exploiters lacked fucoidanases but encoded one of two alternative L-fucose catabolic pathways described in MetaCyc^42^: L-fucose degradation I or L-fucose degradation II. For each pathway, genomes were required to encode at least 75% of the constituent enzymes. For pathway I, this included: K07248 (lactaldehyde dehydrogenase), K02431 (L-fucose mutarotase), K00879 (L-fuculokinase), K01818 (L-fucose/D-arabinose isomerase), and K01628 (L-fuculose-phosphate aldolase). For pathway II, this included: K18333 (L-fucose dehydrogenase), K18334 (L-fuconate dehydratase), K18335 (2-keto-3-deoxy-L-fuconate dehydrogenase), K07046 (L-fuconolactonase), K18336 (2,4-didehydro-3-deoxy-L-rhamnonate hydrolase), and K01685 (altronate hydrolase). Scavengers were defined as genomes lacking all enzymes associated with both fucoidan degradation and L-fucose catabolism.

### Identification of fucoidan polysaccharide-utilization loci (PULs)

We searched closed genomes of cultured fucoidan degraders for candidate polysaccharide utilization loci (PULs) using a sliding 12-gene window. Regions containing at least four carbohydrate-active enzyme (CAZyme) genes were flagged, and those encoding two or more known fucoidanases (GH29, GH95, GH141, GH107, GH168, S1_15, S1_16, S1_17, S1_22, or S1_25) were retained as putative fucoidan PULs. To avoid misclassification, we manually curated these loci and removed those likely involved in the degradation of other polysaccharides—specifically, five loci from *Flavobacteriia* containing carrageenanases (GH16, GH82, GH150, GH167). The final curated set comprised 35 fucoidan-associated PULs spanning 54 CAZyme families (Supplementary Table 3).

### Classification of enzymes

Enzymes from the fucoidanase families GH29, GH95, GH141, GH107, GH168, S1_15, S1_16, S1_17, S1_22 and S1_25, which target sulfated fucose, were classified as ‘backbone’ enzymes. The remaining 44 CAZyme families present in fucoidan PULs were classified as ‘debranching’ enzymes. Predicted biochemical functions of debranching enzymes, based on dbCAN^56^ annotations, included glucuronidases (GH115), mannosidases (GH92, GH130), xylosidases (GH39, GH120), and galactosidases (GH36, GH97, GH117), which may cleave rare monomers in fucoidan branches.

### Definition of total enzyme repertoires

For each degrader, the fucoidanase repertoire was defined as (i) all CAZymes and sulfatases within fucoidan PULs, and (ii) additional chromosomal genes belonging to backbone or rare-monomer families (GH115, GH92, GH130, GH39, GH120, GH36, GH97, GH117). For the characterized degrader^3^, *Lentimonas* sp. CC4, only enzymes that were upregulated at the protein level during growth on *Fucus vesiculosus* fucoidan were included, specifically those in co-expression clusters 2–6.

### Comparison of enzyme repertoires

To sort the sequence diversity into homologous groups of enzymes, we clustered all fucoidan-associated CAZymes using MMseqs2 v13.45111 (easy-cluster with --min-seq-id 0.6 -c 0.5 --cov-mode 0)^58^. This defined enzyme homologs at a threshold of ≥ 60% amino acid identity over ≥ 50% alignment coverage. These clusters formed the basis for comparing enzyme repertoires across isolates, and pairwise similarities were calculated using the Jaccard index.

To explore how enzymatic diversity scales with community size, we performed a rarefaction analysis. In each of 10,000 iterations, we randomly selected *n* degrader strains (*n* = 1–29), counted the number of unique enzyme clusters, and estimated the expected total richness using the Chao1 estimator.

### Identification of primary degraders

To test their ability to grow on fucoidan, all 29 isolates of the strain collection were revived from glycerol stock in 3 mL MBL medium in plastic culture tubes for 3–6 days with carbon sources tailored to their metabolic requirements. Verrucomicrobiota degraders were routinely grown with 0.2% (w/v) fucoidan from *Fucus vesiculosus*. Other degraders (*Flavobacteriia* and Gammaproteobacteria) and exploiters were grown on a mix of 0.2% (w/v) L-fucose 0.2% (w/v) fucoidan from *Fucus vesiculosus*. Scavengers were grown with a mixture of carbon sources matching the isolation medium, each at 0.02% (w/v). Revived cultures were used to inoculate MBL medium supplemented with 0.2% (w/v) fucoidan for 5 days with n = 3 biological replicates for all strains. Growth was monitored by sampling 100 μL of culture and OD_600_ measurements using a Tecan Sunrise plate reader. At the end of growth in early stationary phase, cell-free supernatants were generated by centrifugation at 2200 rpm for 10 min.

### Acid hydrolysis of fucoidan in supernatants

To quantify the decrease of fucoidan-bound monomers after growth, acid hydrolysis was used to cleave the glycosidic linkages of fucoidan remaining in culture supernatants releasing its monomer constituents. Sampled supernatants (5 μl) were mixed with 45 μL of ddH_2_O and 50 μL of 2 M HCl. The HCl solution contained D-galactose-¹³C₆ and D-mannose-¹³C₆ at 15 μM each that were later used as ‘processing internal standards’ to correct technical variability introduced by the sample processing workflow. PCR plates with samples were sealed with plastic strips (Thermo Fisher Scientific AB0600 and AB0784). Acid hydrolysis was carried out for 24h at 100°C in an oven using a custom clamping device to prevent leakage from plates. After hydrolysis, samples were neutralized by addition of 4 M NaOH.

### Derivatization of monosaccharides with 1-phenyl-3-methyl-5-pyrazolone (PMP)

Acid hydrolysates (10 μL sample and 15 μL ddH_2_O) or free monosaccharide samples (25 μL sample) were derivatized with 75 μL of 0.1M PMP in 2:1 methanol:ddH_2_O with 0.4 M ammonium hydroxide for 100 minutes at 70°C following a previously published protocol^37^. For absolute quantification, we used an external calibration curve of a standard mix containing glucuronic acid, xylose, fucose, galactose, mannose ranging from 1 mM to 200 nM prepared in matrices identical to samples. After derivatization, samples and standards were neutralized with 2 M HCl and diluted 1:50 in 0.1% (v/v) formic acid in ddH_2_O containing 50 nM of ‘injection internal standards’, which was used to correct variations in ionization efficiency caused by the ion source of the mass spectrometer. These internal standards consisted of a mix of glucuronic acid, xylose, fucose, galactose, mannose derivatized with heavy labelled PMP-[d5] yielding unique masses distinct from un-labelled PMP.

### Targeted acquisition of PMP-derivatives with LC-MS

PMP-derivatives were measured on a SCIEX qTRAP5500 and an Agilent 1290 Infinity II LC system. The system was equipped with a Waters CORTECS UPLC C18 Column, 90Å, 1.6 μm, 2.1 mm X 50 mm reversed phase column with guard column and 0.2 μM inline filter. The mobile phase consisted of buffer A with 10 mM NH_4_Formate in ddH_2_O and 0.1% (v/v) formic acid and buffer B with 100% acetonitrile and 0.1% (v/v) formic acid. PMP-derivatives were separated by an initial isocratic flow of 13% buffer B for 30 seconds, followed by a binary gradient from 13 % to 36% Buffer B over 2 minutes, followed by a 30 second wash step with 100% buffer B and 30 seconds re-equilibration. The flow rate was constant at 0.5 mL/min with a constant pressure around 430 bar. A diverter valve was used to redirect the first 1.5 minutes of chromatography to waste. The temperature of the column compartment was maintained at 40°C and the autosampler was cooled to 10°C. The ESI source settings were 625°C, with curtain gas set to 30, collision gas to medium, ion spray voltage 5500 and ion source gas 1 and 2 to 90 (arbitrary units).

Data was acquired using multiple reaction monitoring with previously optimized transitions and collision energies in positive mode^36^. For example, a galactose derivative has an exact Q1 mass of 511.2 m/z and was fragmented with a collision energy of 35V to yield the quantifier ion of 175 m/z and the diagnostic fragment of 217.2 m/z. All MRMs and retention times used to identify compounds are listed in Supplementary Table 4.

Typically, each run included 150–250 samples, with randomized injection order. For quality control (QC) samples, we used the highest concentrated standard mix. QC samples and a water blank were injected every 15 samples to monitor consistency and carryover. Calibration curves were prepared in triplicate, and each sample was analyzed in technical duplicates. Guard columns and inline filters were replaced every 500 to 1000 injections.

### Absolute quantification of monosaccharides

Chromatographic data were analyzed using Skyline^59^ v24.1.0.414, with peak areas integrated using default settings and exported for downstream quantification in Python. A reproducible example workflow is provided under https://github.com/EnvSysMicroLAB/Sichert2025_Fucoidan. Briefly, peak areas of target compounds and ^13^C-labeled processing standards were first corrected using the injection internal standards. Subsequently, the corrected ^13^C processing standards were used to normalize the target compound signals. Absolute concentrations of monomers were determined by linear regression against external calibration curves, and technical duplicates were averaged to obtain final concentrations.

### Quantification of monomer degradation

To quantify fucoidan degradation at the monomer level, the concentrations of the five constituent sugars—fucose, glucuronic acid, galactose, mannose, and xylose—in culture supernatants and were compared to uninoculated control media processed in parallel. Notably, the concentrations of ‘rare’ monomers are calculated from the sum of the four non-fucose sugars, while ‘total” monomers represent the sum of all five monomers. Degradation *f* of a given monosaccharide *i* (*i* ∈{Fuc, GlcA, Gal, Man, Xyl, Rare, Total}) by strain or strain combination *j,* was quantified as:

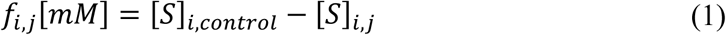

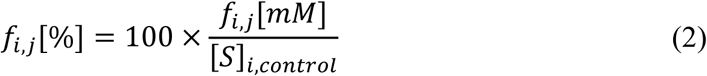

where [S]_i,control_ is the concentration in the uninoculated control, and [S]_i,j_ is the concentration after growth of strain *j*.

For every strain–monomer combination, we tested whether the observed decrease in concentration exceeded abiotic background using a one-sided t-test (alternative = “less”). Resulting p-values were corrected for multiple comparisons using the Benjamini–Hochberg false-discovery-rate procedure (FDR, α = 0.01). Degradation was considered significant when both the raw p-values and the FDR-adjusted q were < 0.01. Throughout this study, we generated five distinct datasets, with monosaccharide concentrations and corresponding statistical analyses reported in Supplementary Table 5. Unless otherwise stated, all statistical analyses of degradation data involving Pearson correlation coefficients and associated P-values were performed using replicate-level data from three biological replicates, without prior aggregation into means.

### High-throughput community experiments

To assess the impact of biological interactions on fucoidan degradation, we performed four high-throughput growth experiments using a standardized cultivation setup. All experiments were conducted in biological triplicates (n = 3) at 20 °C with orbital shaking at 200 rpm. Glycerol stocks were revived in 3 mL Marine Broth Lite (MBL) in plastic culture tubes for 3– 6 days, using carbon sources matched to the metabolic requirements described above. Revived cultures were diluted 1:5 into fresh medium in 96-deep-well plates (1.8 mL per well), each containing a 4 mm glass bead (Sigma-Aldrich, Z143936), and incubated for 18 h to reach exponential phase (OD₆₀₀ = 0.15–0.3). Exponential-phase cells were washed in carbon-free MBL and used to inoculate experimental cultures. For monocultures, the initial OD₆₀₀ was set to 0.01. Co-cultures were started by adding each strain at OD₆₀₀ = 0.01, as detailed below. Experimental cultures were incubated for up to 5 days in the same deep-well plate format, sealed with breathable lids (Kuhner, 104098 and 104106). At defined time points, 100 μL of culture was sampled for OD₆₀₀ measurement using a Tecan Sunrise plate reader. Samples were centrifuged at 2200 rpm for 10 min to separate cells and supernatant; supernatants were stored at –20 °C for further analysis.

### Verrucomicrobiota-centric pairwise co-cultures

To assess the influence of community members on the degradation activity of primary degraders, each of the 29 isolated strains was co-cultured in a 1:1 ratio (normalized by OD₆₀₀) with one of the three primary degraders (V4, V25, or V69), yielding 87 distinct pairwise combinations. Cultures were incubated for 5 days, and fucoidan degradation was quantified at the final time point. Growth dynamics in co-cultures were assessed using two complementary metrics: (i) total biomass, defined as the mean of the two highest OD₆₀₀ values recorded during the time course, and (ii) apparent growth rate, calculated as the inverse of the time required to reach an OD₆₀₀ threshold of 0.15 (1/tₜₕ). In co-cultures of V69 with strains F12, F40, F56, and G88, we observed aggregate formation that interfered with OD₆₀₀-based biomass estimates. For these cases, total biomass was instead determined by quantifying bacterial DNA from 100 μL of culture using the DNAdvance Kit (Beckman Coulter) followed by absolute DNA quantification with the Femto Bacterial DNA Quantification Kit, according to the manufacturer’s protocol.

### Classification of pairwise interactions

To identify potential synergistic or antagonistic effects in pairwise co-cultures, we compared biomass formation and fucoidan degradation to the corresponding monocultures. Differences in biomass were considered significant if both the p-value and the false discovery rate (FDR, Benjamini–Hochberg correction, α = 0.01) were <0.01. Differences in degradation were considered significant if both the p-value and FDR (α = 0.05) were <0.05 and the absolute log₂ fold change exceeded 0.1.

### Synergism beyond resource-partitioning

To assess synergistic interactions in pairwise co-cultures, we formulated a null model that estimates expected degradation outcomes based on monoculture performance, assuming no metabolic complementarity between strains. This model reflects a scenario in which both strains have fully overlapping substrate preferences, such that each monomer can be degraded only up to the level achieved by the better-performing monoculture. The difference between this null expectation and the observed co-culture degradation defines the synergism score σ(i, j), formally given as:

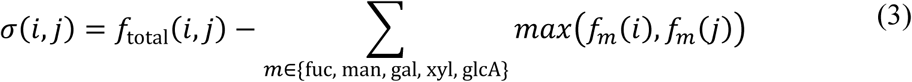

where *f(i, j)* is the total fraction of fucoidan degraded in co-culture of strains *i* and *j*, *f_m_(i)* and *f_m_(j)* are the fractions of monomer m degraded by strains *i* and *j* in monoculture. This conservative formulation avoids overestimation of expected degradation by preventing redundant contributions from metabolically similar strains. Positive values of *σ(i, j)* thus indicate cooperation through functional complementarity, whereas negative values suggest antagonism. To determine whether observed co-culture degradation significantly deviated from the null expectation, we applied a two-sided *t*-test. Resulting *P*-values were corrected for multiple comparisons using the Benjamini–Hochberg procedure, and synergism scores were considered significant if they satisfied *P* < 0.01, *q* < 0.01, and exhibited an absolute effect size greater than 5 (|σ| > 5). Notably, this framework can be extended to communities with more than two members by using the maximum monomer degradation observed in the best-performing drop-out community as the null expectation.

### Cross-feeding of residual fucoidan

Partially degraded fucoidan was recovered from cultures of V4, V25, and V69 grown to late stationary phase in 1 L MBL medium supplemented with 0.2% (w/v) fucoidan from *Fucus vesiculosus*. The sterile filtered supernatant was concentrated using an Amicon stirred ultrafiltration cell (EMD Millipore, UFSC20001) with a 1-kDa cellulose membrane (EMD Millipore, PLAC06210) on ice using nitrogen for gas pressure. Desalting was carried out by addition of ddH_2_O and continued concentration. Concentrated material was lyophilized.

To test how ‘secondary’ degraders could utilize the residuals substrates of primary degraders, F12, F40, F56, F94, V25, V69, and V4 were grown in MBL medium containing 0.1% (w/v) of each residual fucoidan. Cultures were incubated for 4 days, and degradation was quantified from supernatants collected at the final time point compared to initial concentrations.

To compare the additional degradation of residuals achieved by secondary degraders to the additional degradation observed in co-cultures, we used the relationship:

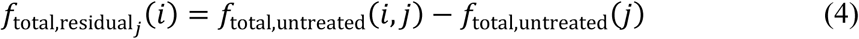

Here, *f_total,residual_(i)* denotes the degradation of secondary degrader *i* on residuals of primary degrader *j*; *f_total,untreated_(i, j)* is the total degradation observed in co-culture of strains *i* and *j* on untreated fucoidan; and *f_total,untreated_(j)* is the degradation by strain *j* alone on the same substrate.

### Full Combinatorial Degrader Communities

To systematically evaluate synergistic interactions in fucoidan degradation, we constructed all 127 possible combinations of the seven degraders F12, F40, F56, F94, V25, V69, and V4, along with a no-cell negative control. Each community was assembled by mixing strains at equal optical density (OD₆₀₀) at 0.01 each and inoculating into 1.8 mL of MBL medium supplemented with 0.2% (w/v) fucoidan from *Fucus vesiculosus* as the sole carbon source. Cultures were incubated for 5 days (n = 3 biological replicates), after which supernatants were collected for degradation analysis.

### Modelling

To predict degradation from community composition, we developed a nonlinear trait-based model in which each bacterial strain contributes independently to the degradation of two classes of fucoidan-derived monomers: fucose and rare sugars. Each strain *i* was assigned two parameters (*w_iFuc_* and *w ^Rare^*) representing its degradation capacity for these monomer types. For a given community composed of *m* strains, represented by a presence/absence vector *x_i_*∈*{0,1}*, the total effective capacity to degrade each monomer pool was computed as the sum of contributions from all present strains:

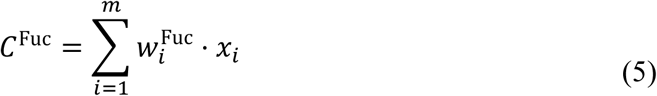

To capture the saturating behavior observed in experimental data, we applied a Hill function to these summed capacities, introducing nonlinearity and a threshold-like response:

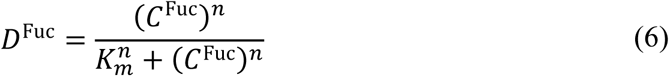

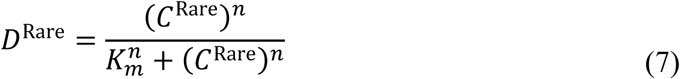

Here, *K_m_^n^* is the half-saturation constant, and *n* is the Hill coefficient, both shared across the two monomer classes. The resulting values *D_Fuc_* and *D_Rare_* represent the predicted fraction of each monomer pool degraded by the community.

Experimental degradation values of 127 communities were used to fit the model parameters: one degradation capacity per strain and per monomer pool (*w_iFuc_* and *w_iRare_*), and shared parameters *K_m_^n^* and *n*. Model parameters were inferred by minimizing the root mean squared deviation between predicted and observed degradation. Optimization was performed in Julia using the LsqFit.jl package. Initial and final parameters values, as well as their effect on model fit are shown in Extended Data Fig. 9a-c. Goodness-of-fit was assessed via visual comparison and R^2^ statistics (Extended Data Fig. 9d). All code and data used in model construction and fitting is available here https://github.com/EnvSysMicroLAB/Sichert2025_Fucoidan.

### Degradation of diverse fucoidans

To evaluate the degradation capabilities of individual degraders across diverse fucoidans, we assembled a panel of nine brown algal fucoidans. These included fucoidans derived from seven brown algal species selected for their pronounced differences in composition and structure, as well as three variants of *Fucus vesiculosus* sourced from three different vendors. The latter were included to capture subtle compositional and structural variations suspected to arise from differences in sampling time, sampling season, and extraction method^27,30^. We tested the growth of all seven degrader strains in monoculture on each of these nine fucoidans.

For community experiments, degraders were grouped to minimize redundant combinations and maximize between-group complementarity. Group A comprised strains V25, V4, and F56. Group B included strains F12, F40, and F94. Group C consisted of strain V69. These groups were tested individually, as well as in all pairwise and triplet combinations, resulting in seven unique community configurations. Each community was inoculated with equal OD₆₀₀ contributions from each strain into 1.8 mL MBL medium containing 0.2% (w/v) of one of the nine fucoidans. Cultures were incubated for 5 days (n = 3), and degradation was quantified as described above.

These experimental data—63 total community–substrate combinations—were used to validate the model with out-of-sample predictions. To account for variability in monomer composition across polymers, we weighted the contribution of each monomer type to total degradation according to its relative abundance in the polymer.

### Reporting summary

Further information on research design is available in the Nature Portfolio Reporting Summary linked to this article.

## Data availability

All data supporting the findings of this study are available within the paper and its Supplementary Information. Accession numbers for all genomes used in this study are listed in Supplementary Table 2. Raw LC-MS data are available via Panorama Public at https://doi.org/10.6069/r2z3-7b10. Genome assemblies and associated sequencing data have been deposited in the NCBI Sequence Read Archive under accession number PRJNA878624.

## Code availability

A reproducible workflow for calculating degradation from LC-MS data, along with code for model implementation and validation, is available at: https://github.com/EnvSysMicroLAB/Sichert2025_Fucoidan.

## Acknowledgments

We thank Stuart Levine for providing sequencing infrastructure; Angela Kosturanova for technical assistance with DNA extractions and measurements; and Chris Springate for generously providing fucoidan from *Laminaria hyperborea*. We are grateful to Bram Vekeman for sharing his expertise in bacterial isolation, Alaa Othman and Nicola Zamboni for providing metabolomics expertise and infrastructure, and Sammy Pontrelli and Julia Schwartzman for stimulating discussions. This project was supported by the Simons Foundation through the Principles of Microbial Ecosystems (PRIME) collaboration (grant 542395). A.S. was supported by the European Molecular Biology Organization Postdoctoral Fellowship (ALTF Grant ALTF 996-2021). A.G. acknowledges support from the Ashok and Gita Vaish Junior Researcher Award, the DST-SERB Ramanujan Fellowship, as well the DAE, Govt. of India, under project no. RTI4001.

## Author contributions

A.S. and O.X.C. conceived the study. All authors contributed to the experimental design. A.S. performed the investigations and analysed the data. S.P. and A.G. designed and implemented the modelling framework. A.S., O.X.C. and U.S. wrote the manuscript with input from all authors.

## Competing interests

The authors declare no competing interests.

## Extended Data Figures

**Extended Data Fig. 1:**
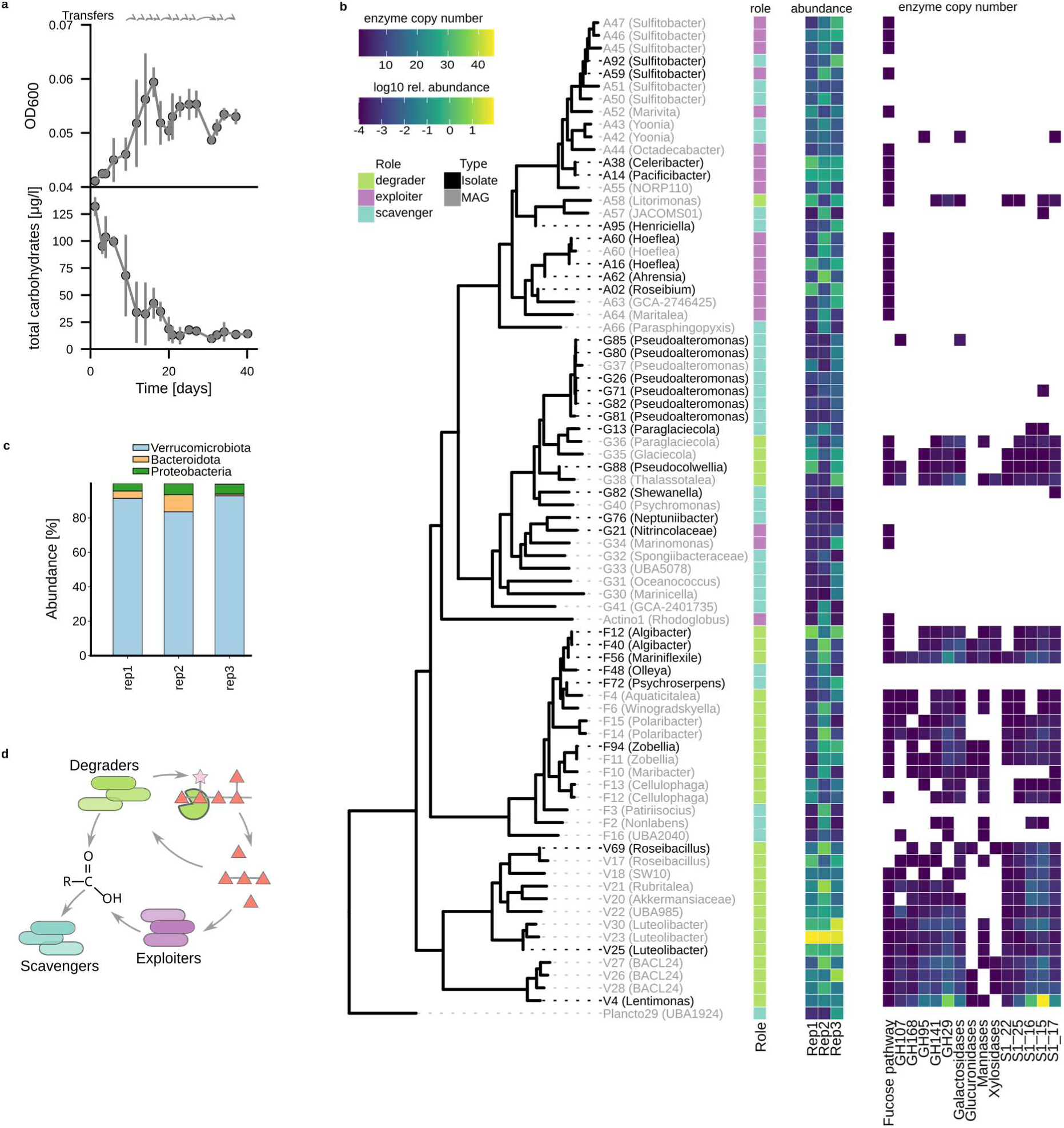
Genome centric analysis of fucoidan enrichments. **a**, Time course of growth and total carbohydrate concentrations; data points and error bars represent mean ± s.d. (n = 3), grey arrows indicate 1:50 transfers into fresh medium. **b,** Maximum-likelihood phylogeny of MAGs and isolates reconstructed from the enrichments, based on a concatenated alignment of 120 single-copy marker genes. Accompanying heatmaps show (from left to right): (i) inferred metabolic roles (e.g., degrader, exploiter); (ii) relative abundance at the final time point in all three replicates; (iii) presence of a complete fucose degradation pathway and counts of homologs in selected sulfatase and CAZyme families. **c,** Phylum-level composition at the final time point. **d,** Schematic of metabolic interactions in fucoidan-degrading communities. Degraders initiate fucoidan degradation via extracellular enzymes. Breakdown products are utilized by both degraders and exploiters, while scavengers rely on metabolic waste products released by degraders or exploiters.

**Extended Data Fig. 2:**
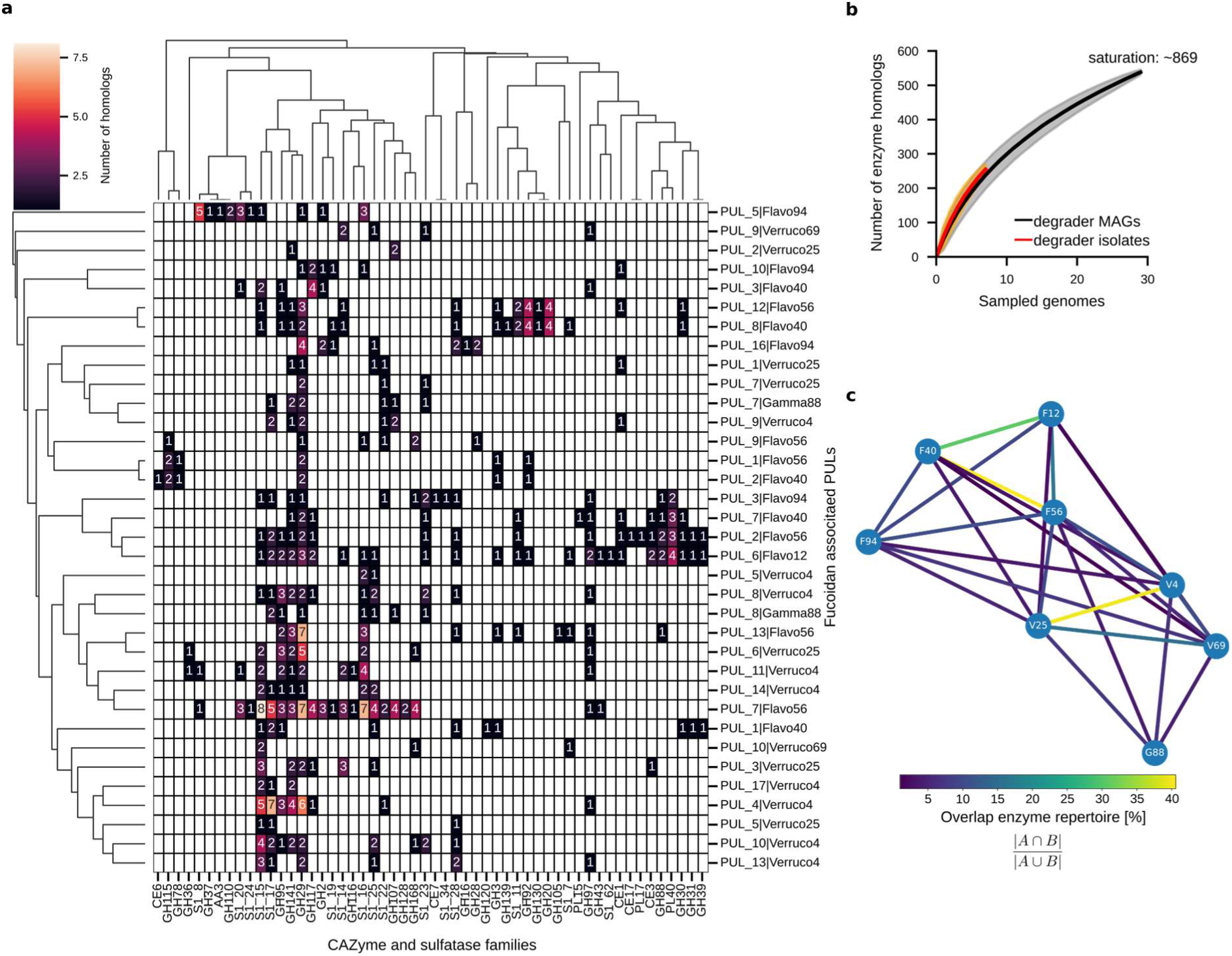
Comparison of enzyme repertoires in fucoidan-degrading genomes. **a**, Heatmap of homolog counts per enzyme family across polysaccharide utilization loci (PULs) in isolated degraders associated with fucoidan degradation. Each column corresponds to a PUL, each row to an enzyme family. Numbers denote the count of homologs per family per locus. Dendrograms reflect hierarchical clustering based on pairwise correlation. Known fucoidanases used to identify PULs are highlighted in red. **b**, Rarefaction analysis of enzyme homologs identified in isolated degraders (orange) and in both isolates and MAGs (black). Solid lines indicate the average number of unique homologs per genome; shaded areas show s.d. from 10,000 random samplings. Saturation was estimated using the Chao1 estimator. **c,** The nodes of the network represent isolated degraders and the edges are coloured by the normalized similarity of their enzyme repertoires calculated from the Jaccard similarity.

**Extended Data Fig. 3:**
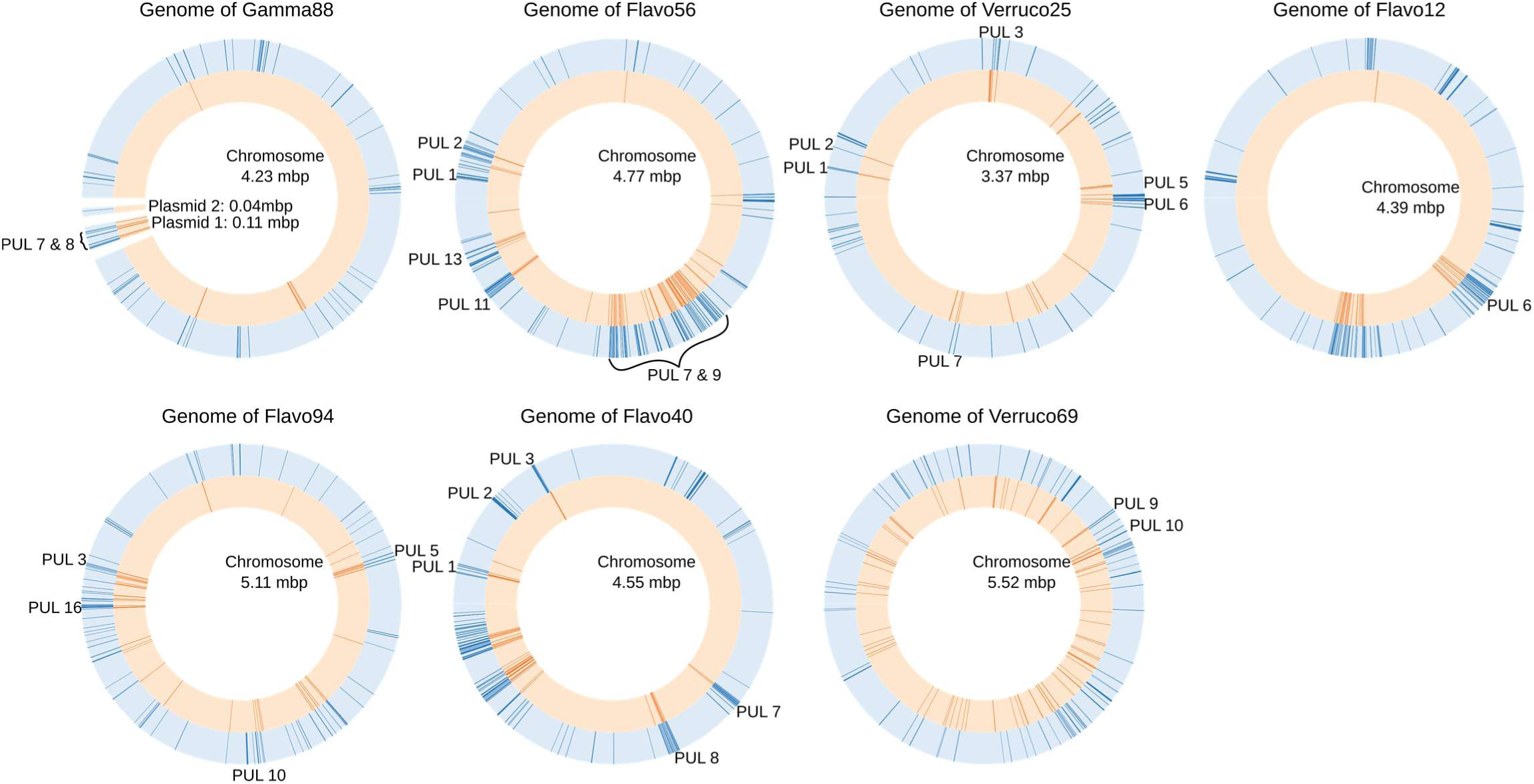
Genomic localization of fucoidan-degrading polysaccharide utilization loci. Genomic context of all seven newly isolated fucoidan-degrading strains used in this study. Circular genome plots show the distribution of carbohydrate-active enzymes (blue) and sulfatases (orange). Labelled regions denote polysaccharide utilization loci associated with fucoidan degradation.

**Extended Data Fig. 4:**
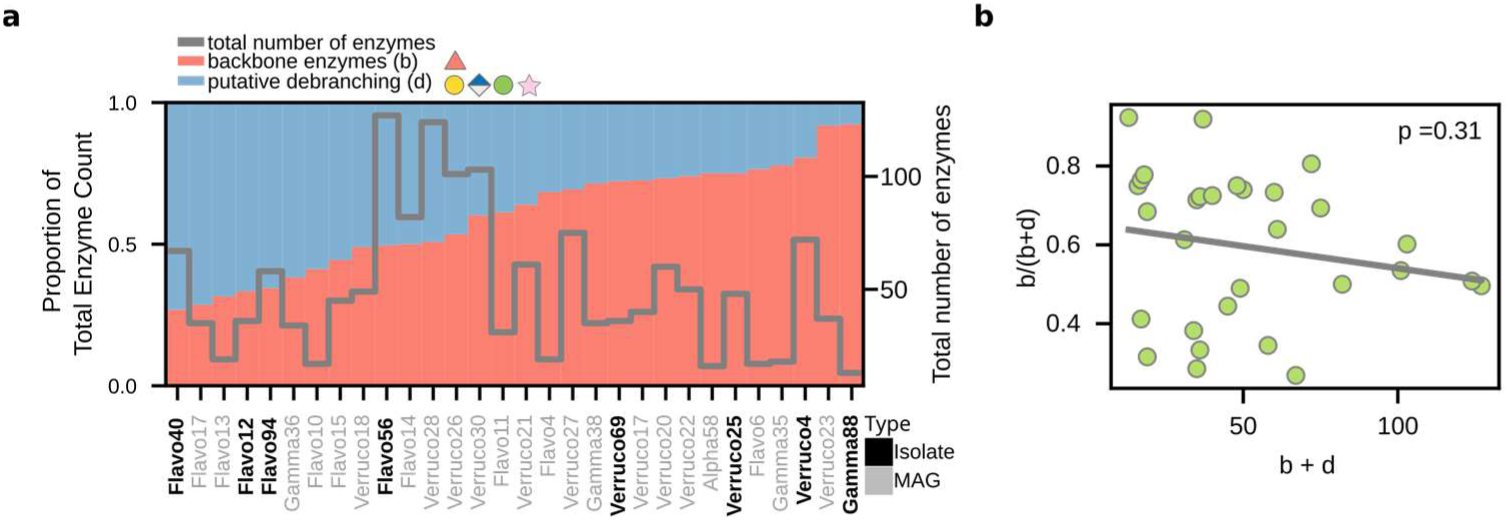
Enzymatic specialization of fucoidan degraders. **a**, Proportion of backbone-targeting (blue) and putative debranching (red) enzymes in each fucoidan PUL, across genomes of isolated (black) and metagenome-assembled (grey) degraders. Backbone-targeting enzymes include all characterized enzymes acting on sulfated fucose linkages; debranching enzymes comprise all other CAZyme families co-localized within fucoidan PULs. **b,** Relationship between total number of fucoidan-targeting enzymes and the proportion of backbone-targeting enzymes across 28 degrader genomes. Each point represents one genome; the grey line shows a linear regression (ordinary least squares) with corresponding p-value.

**Extended Data Fig. 5:**
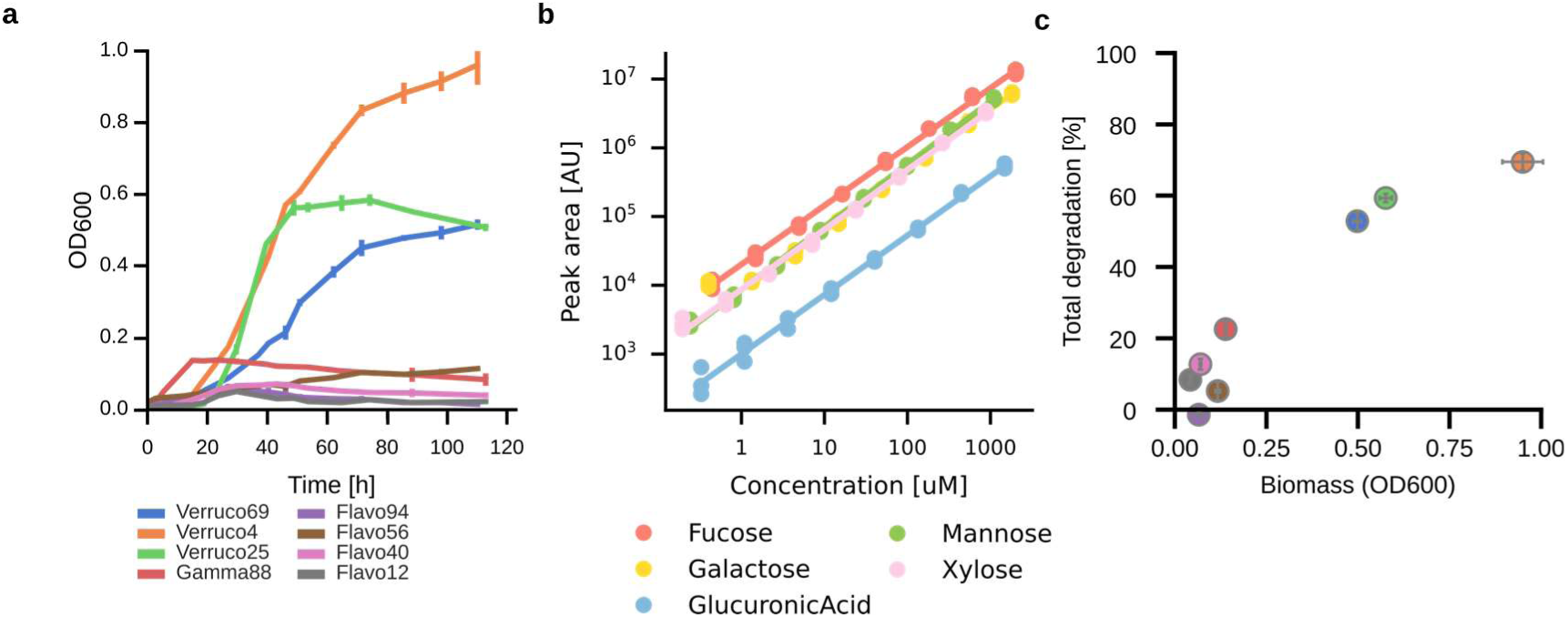
Growth and LC-MS–based quantification of fucoidan degradation. **a**, Growth curves of eight degraders on fucoidan as sole carbon source. Points and error bars show mean ± s.d. of three biological replicates. **b**, Calibration curves for five fucoidan-derived monosaccharides—fucose, galactose, glucuronic acid, mannose, and xylose—used for external quantification. Points represent triplicates of serial dilutions of standards, color-coded by monomer. **c**, Relationship between biomass yield and total fucoidan degradation across eight isolated degrader strains. Data represent mean ± s.d. from three biological replicates.

**Extended Data Fig. 6:**
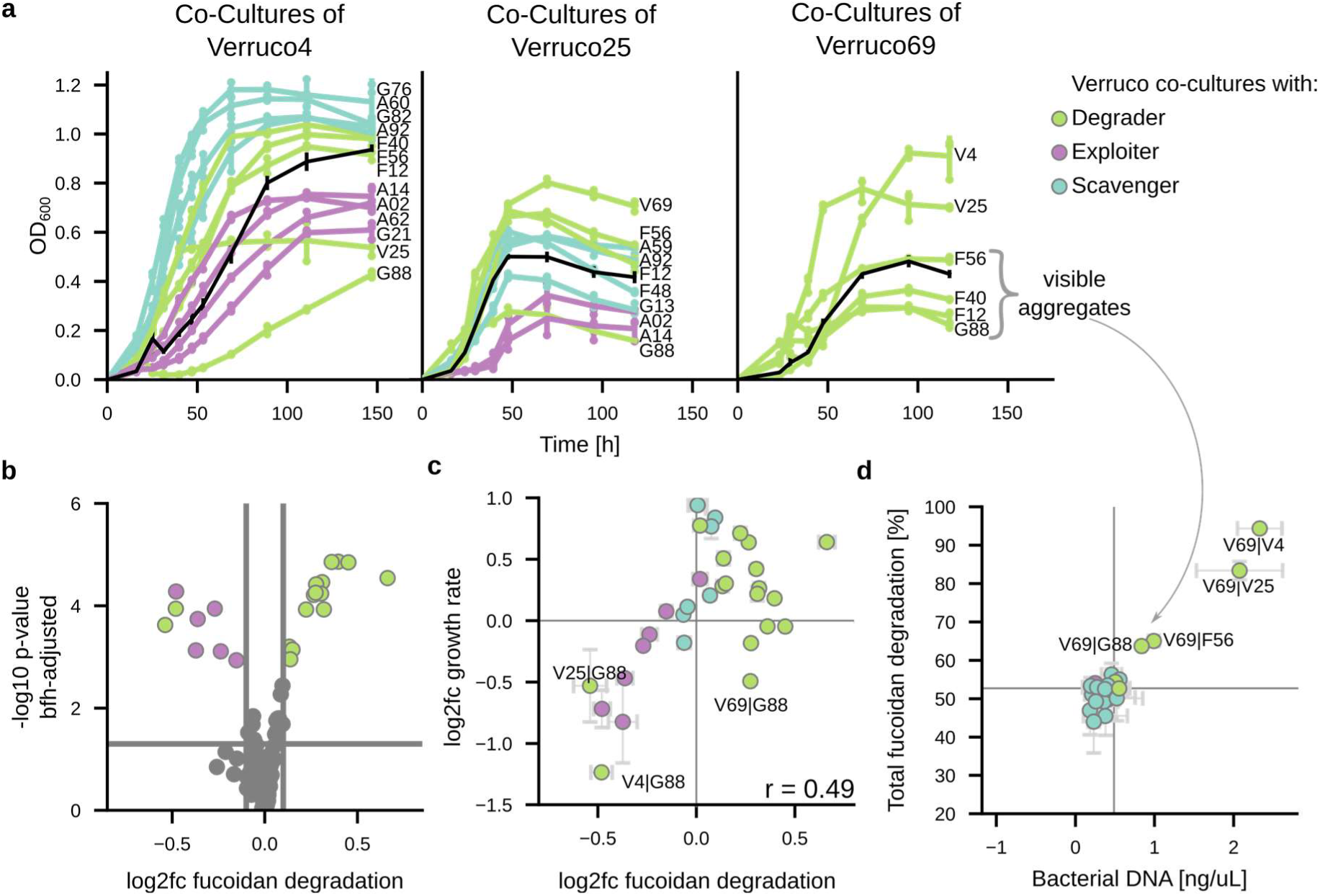
Analysis of growth and fucoidan degradation in Verrucomicrobia-centric co-cultures. **a**, Growth curves of Verrucomicrobia primary degraders (black lines) in pairwise co-culture with partners of different functional roles. Only co-cultures showing significant changes in maximal OD_600_ are displayed; interaction partners are labelled and color-coded by functional role. **b,** Volcano plot showing log_2_-fold changes in fucoidan degradation in co-cultures compared to corresponding monocultures. Coloured points indicate significant changes, grouped by the role of the interaction partner. **c,** Relationship between changes in fucoidan degradation and growth rate across all co-cultures. Only significant differences are shown. **d,** Comparison of total bacterial biomass (measured by qPCR targeting bacterial DNA) and fucoidan degradation across all co-cultures of Verruco69. The grey arrow highlights four co-cultures (with G88, F56, F12, and F40) that formed visible precipitates, which interfered with optical density (OD_600_) measurements shown in panel **a**. For these four co-cultures, OD-based estimates of changes in biomass were substituted by DNA-based estimates, including those shown in Fig. 2a. All data points and error bars represent mean ± s.d. of three biological replicates.

**Extended Data Fig. 7:**
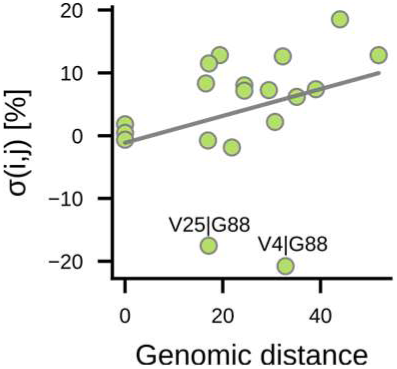
Synergism scores in degrader co-cultures as a function of genomic distance. Pairwise synergism scores (σ(i,j)) plotted against the Euclidean distance in the proportion of backbone-targeting enzymes relative to total fucoidan-associated enzymes. Each data point represents the mean synergism score from three biological replicates; the grey line indicates a linear regression. Selected strain pairs are labelled. The co-cultures V4|G88 and V25|G88 are included in the analysis but fall outside the main trend.

**Extended Data Fig. 8:**
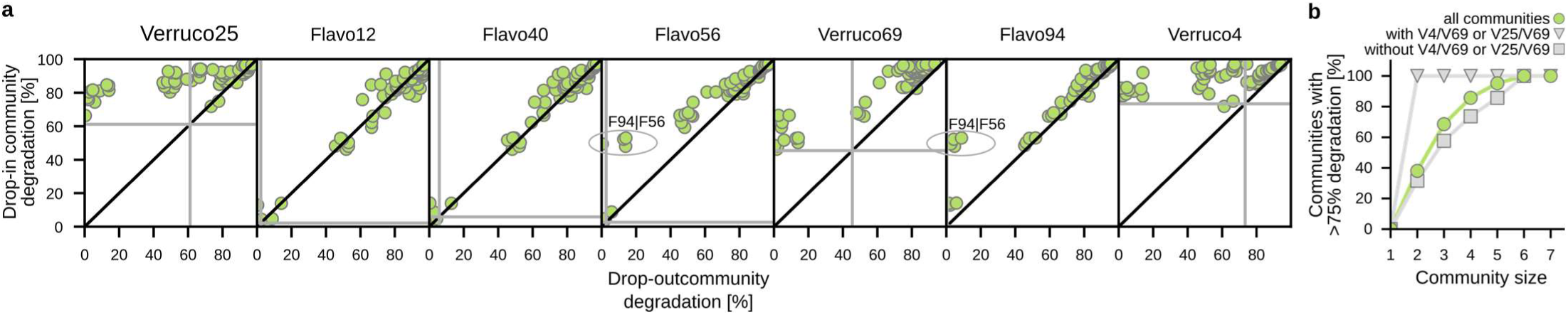
Fucoidan degradation in seven member communities. **a**, Each panel shows a pairwise comparison of total fucoidan degradation in communities with (y-axis) versus without (x-axis) a given focal strain, across all 63 unique community combinations containing that strain. The black diagonal line indicates equal degradation in both conditions. Grey horizontal lines indicate the degradation achieved by the focal strain in monoculture. Circled examples highlight strong emergent interactions between strains F56 and F94. **b**, Percentage of communities achieving ≥75% fucoidan degradation, grouped by community size (1 to 7 strains). Green circles represent all communities (n = 127). Grey triangles indicate communities containing the strain pairs V4/V69 or V25/V69; grey squares indicate those without either pair. The 75% threshold was chosen to reflect degradation levels exceeding those of the best-performing monocultures.

**Extended Data Fig. 9:**
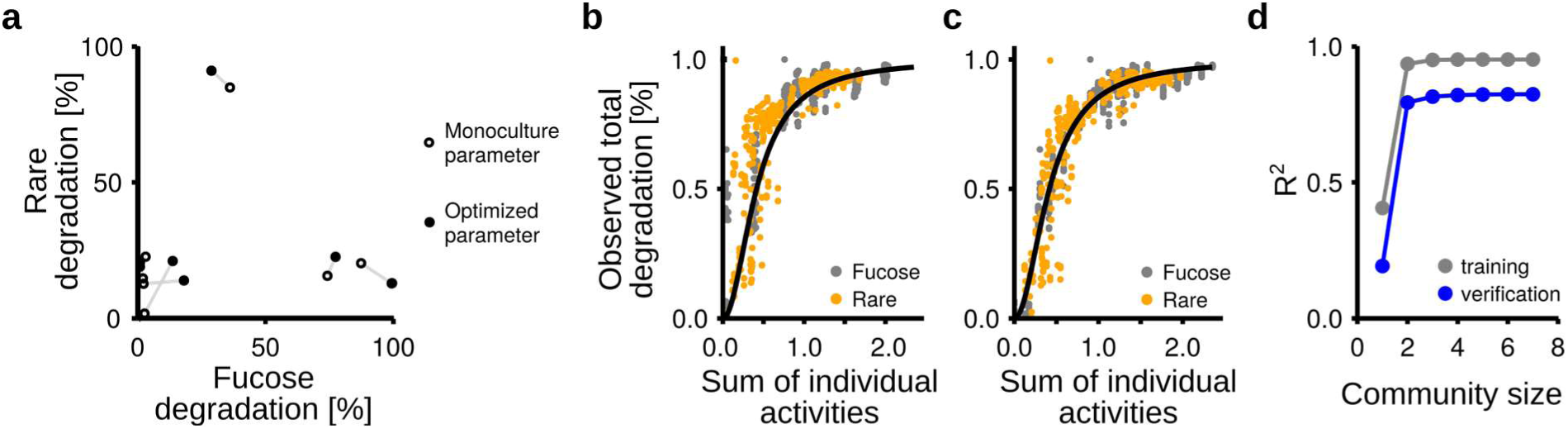
Trait-based modelling of community degradation. **a**, Effect of parameter optimization on the monomer degradation of seven degraders. Circles represent strain-specific degradation capacities before (hollow) and after (filled) optimization. Observed values were averaged from three biological replicates per strain. **b**, Model prediction using original monoculture-derived activities for each monomer pool. The black line shows a fitted Hill function. Grey and orange data points represent fucose and rare monomer degradation, respectively, measured across 127 communities in biological triplicates. **c**, Same as (**b**), but using optimized strain activity parameters. **d**, Model performance (R²) as a function of maximal community size included in training.

**Extended Data Fig. 10:**
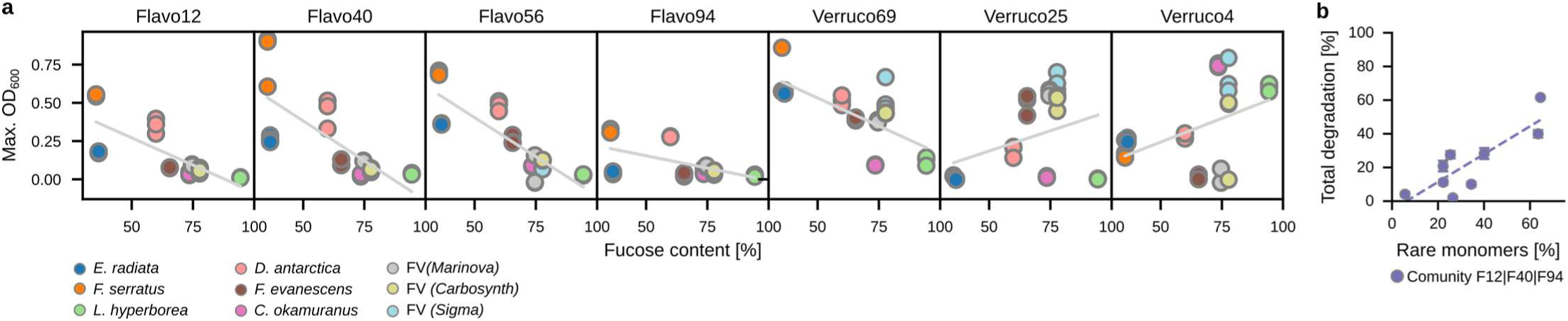
Influence of fucoidan monomer composition on growth and degradation. **a**, Growth of individual strains on different fucoidans, shown as a function of fucose content. All data points from three biological replicate are shown; points are color-coded by the fucoidan substrate. Grey lines indicate linear regressions. **b,** Total degradation of fucoidan by a community of three *Flavobacteriia* as a function of monomer composition. Dashed line indicates a linear regression. Data points and error bars represent the mean ± s.d. of three biological replicates.

## Supplementary Table Legends

**Supplementary Table 1**

Media formulations used in this study.

**Supplementary Table 2**

Sheet 1: Genome identifiers, accession numbers, taxonomy, strain names, and inferred functional roles.

Sheet 2: Relative abundances of genomes in metagenomes of enrichment cultures.

**Supplementary Table 3**

Sheet 1: Overview of enzyme classes and predicted functions.

Sheet 2: Genomic localization of all fucoidan-associated enzymes across genomes. Sheet 3: Protein sequences of all fucoidan-degrading enzymes.

Sheet 4: MMseqs2 clustering of fucoidan-degrading enzymes.

**Supplementary Table 4**

Sheet 1: Exact masses, fragment ions, and retention times used to identify fucoidan monomers by LC-MS.

Sheet 2: Sugar concentrations, calculated degradation, and statistical significance for 29 bacterial strains grown in monoculture.

Sheet 3: Sugar concentrations, calculated degradation, and statistical significance for 87 pairwise co-cultures.

Sheet 4: Sugar concentrations, calculated degradation, and statistical significance for seven degraders grown on residual fucoidans from primary degraders.

Sheet 5: Sugar concentrations, calculated degradation, and statistical significance for all 127 combinations of seven degraders grown on fucoidan.

Sheet 6: Sugar concentrations, calculated degradation, and statistical significance for seven communities grown on nine different fucoidans.

**Supplementary Table 5**

Sheet 1: Growth measurements of 87 pairwise co-cultures.

Sheet 2: Fold-changes and statistical significance of fucoidan degradation in 87 pairwise co-cultures.

Sheet 3: Fold-changes and statistical significance of growth in 87 pairwise co-cultures.

Sheet 4: Fold-changes and statistical significance of total bacterial DNA in 29 pairwise co-cultures involving Verruco69.

Sheet 5: Calculated synergism scores and statistical significance for 87 pairwise co-cultures.

**Supplementary Table 6**

Observed and predicted degradation values used for model training, testing, and validation.

## Notes

### Competing Interest Statement

The authors have declared no competing interest.

