## Supplementary figures and images for "A division of labor controls the degradation of fucoidans in the ocean"

### ED1.png

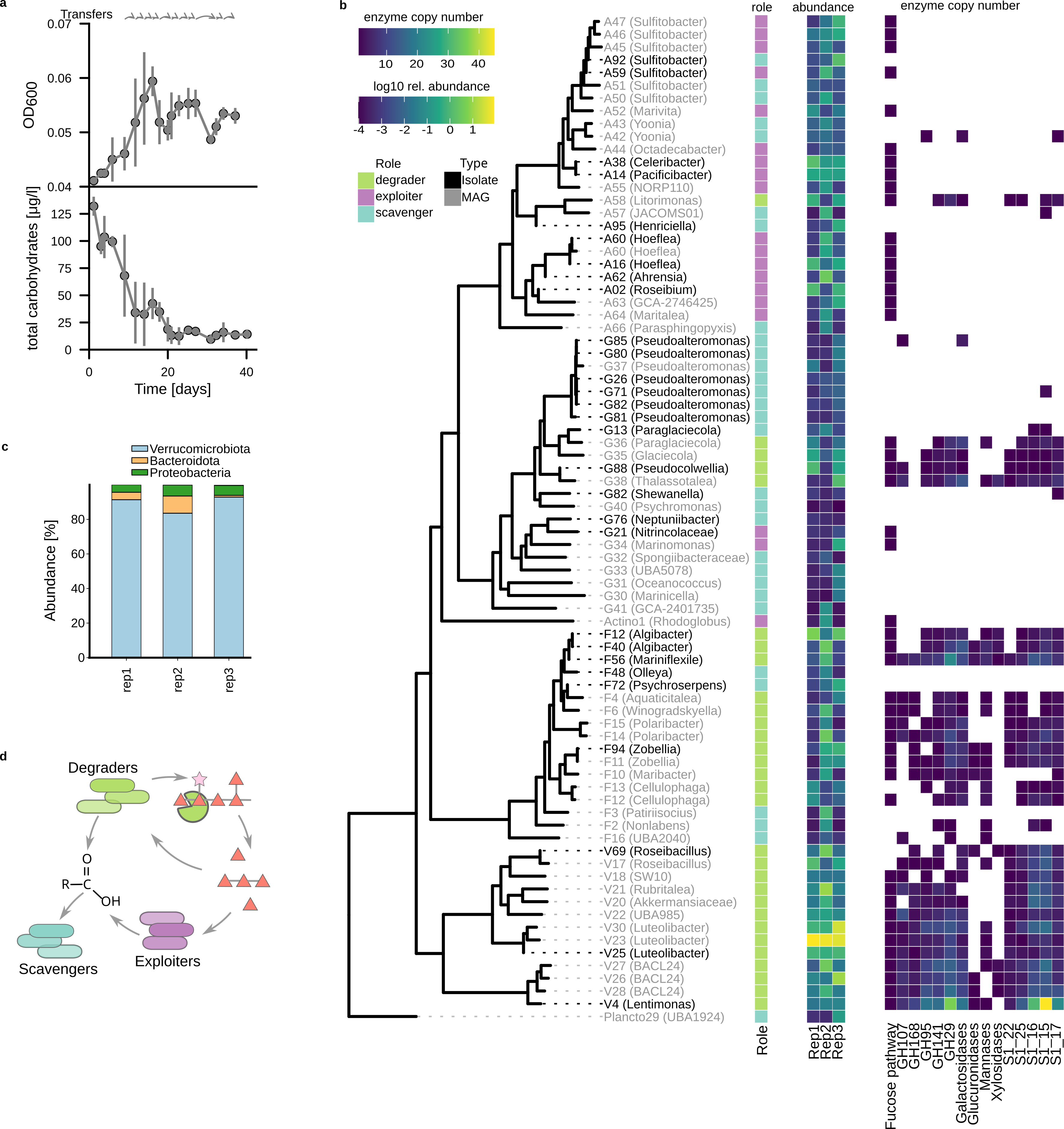

### ED2.png

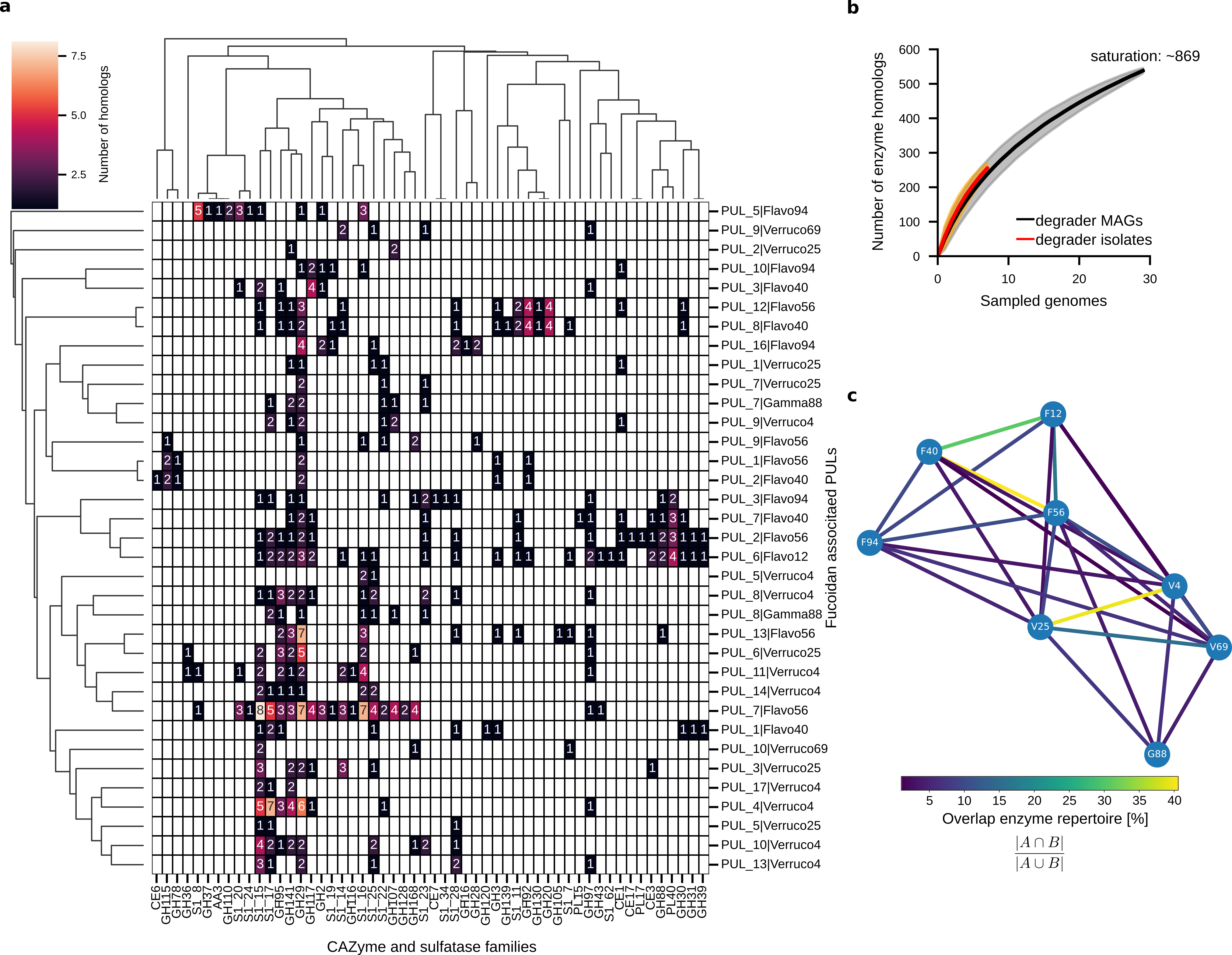

### ED3.png

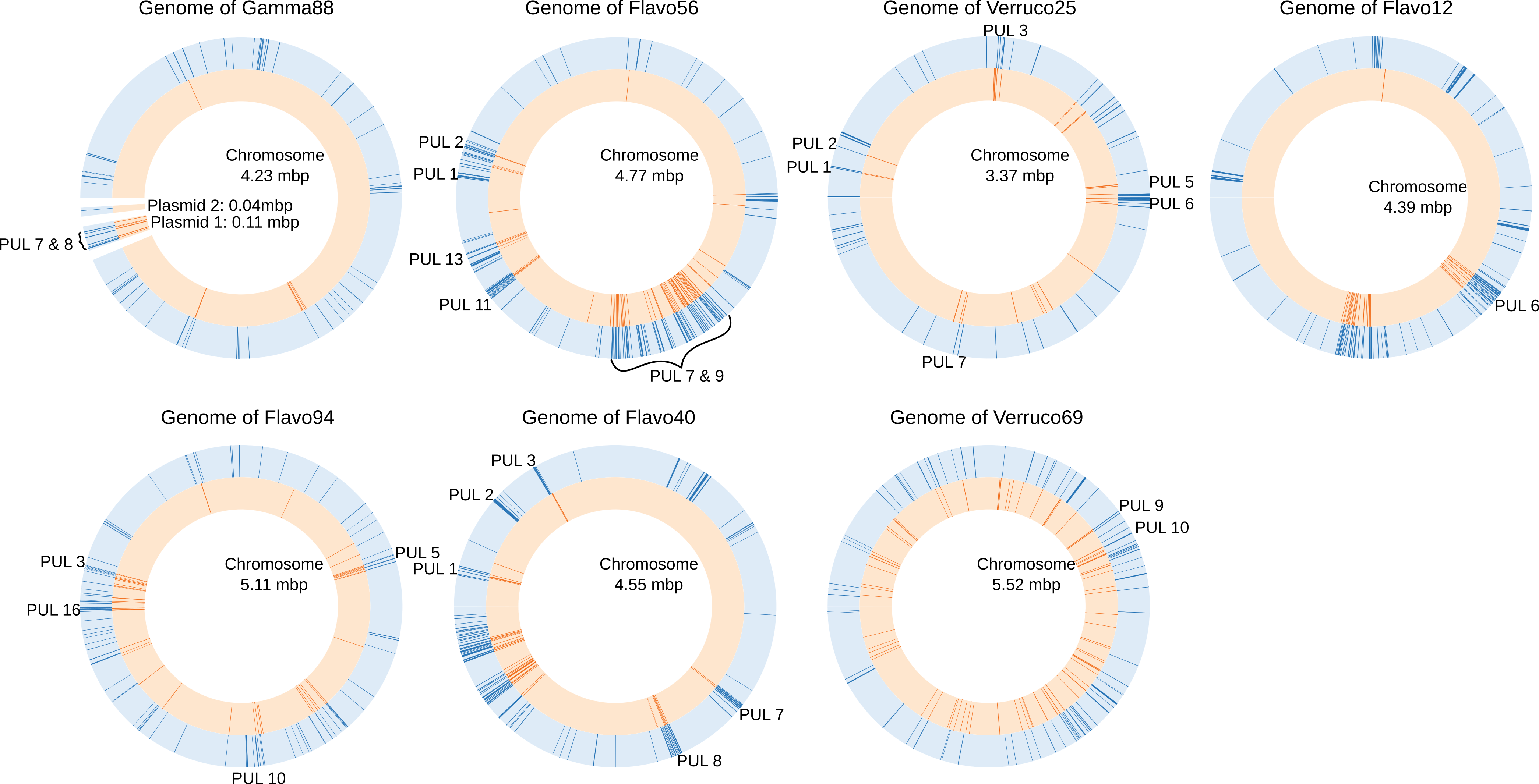

### ED4.png

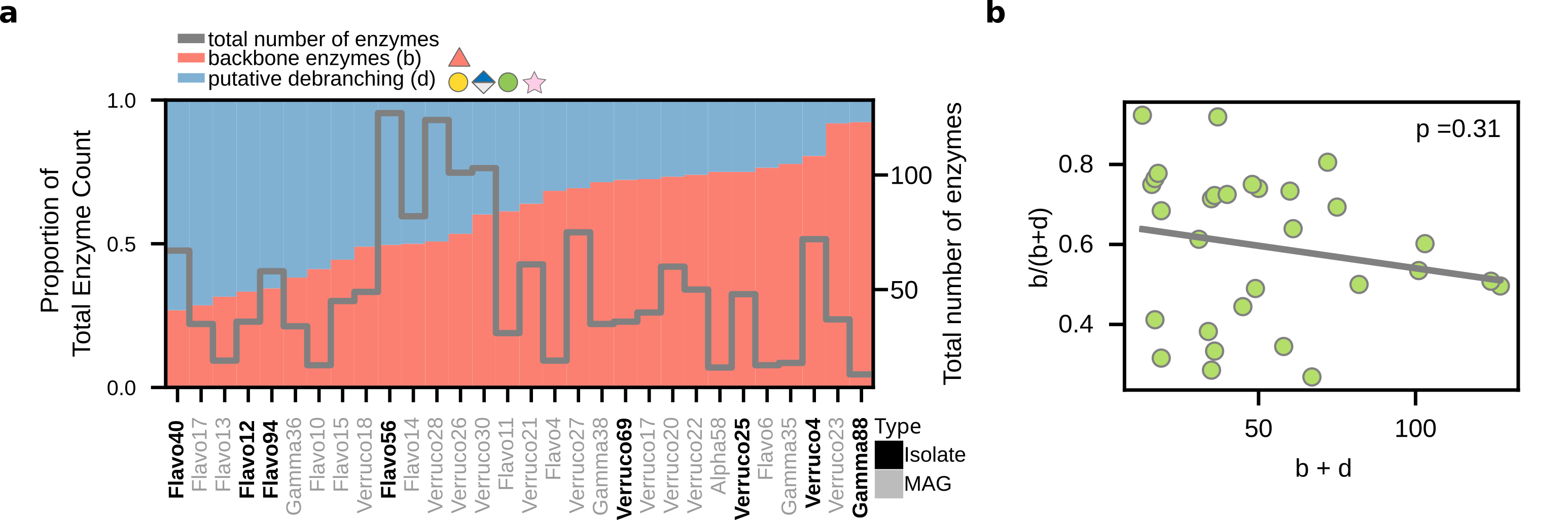

### ED5_qTRAP.png

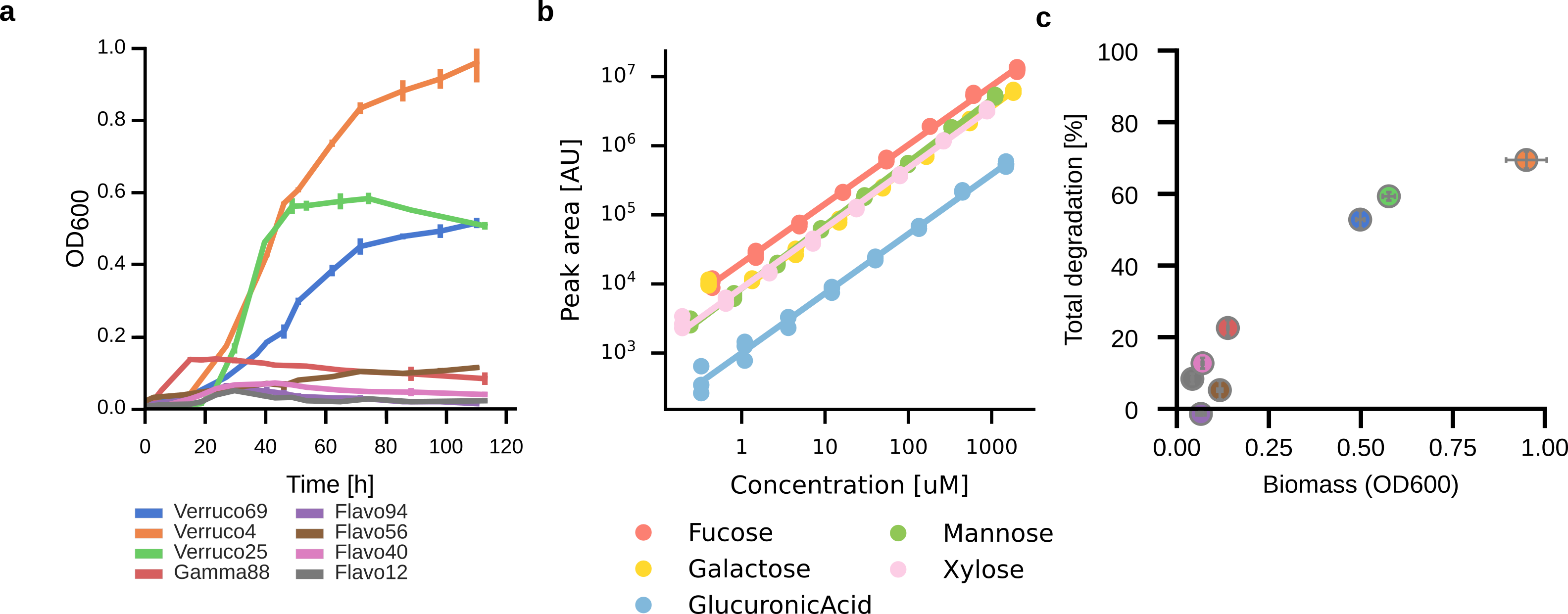

### ED6_CoCultures.png

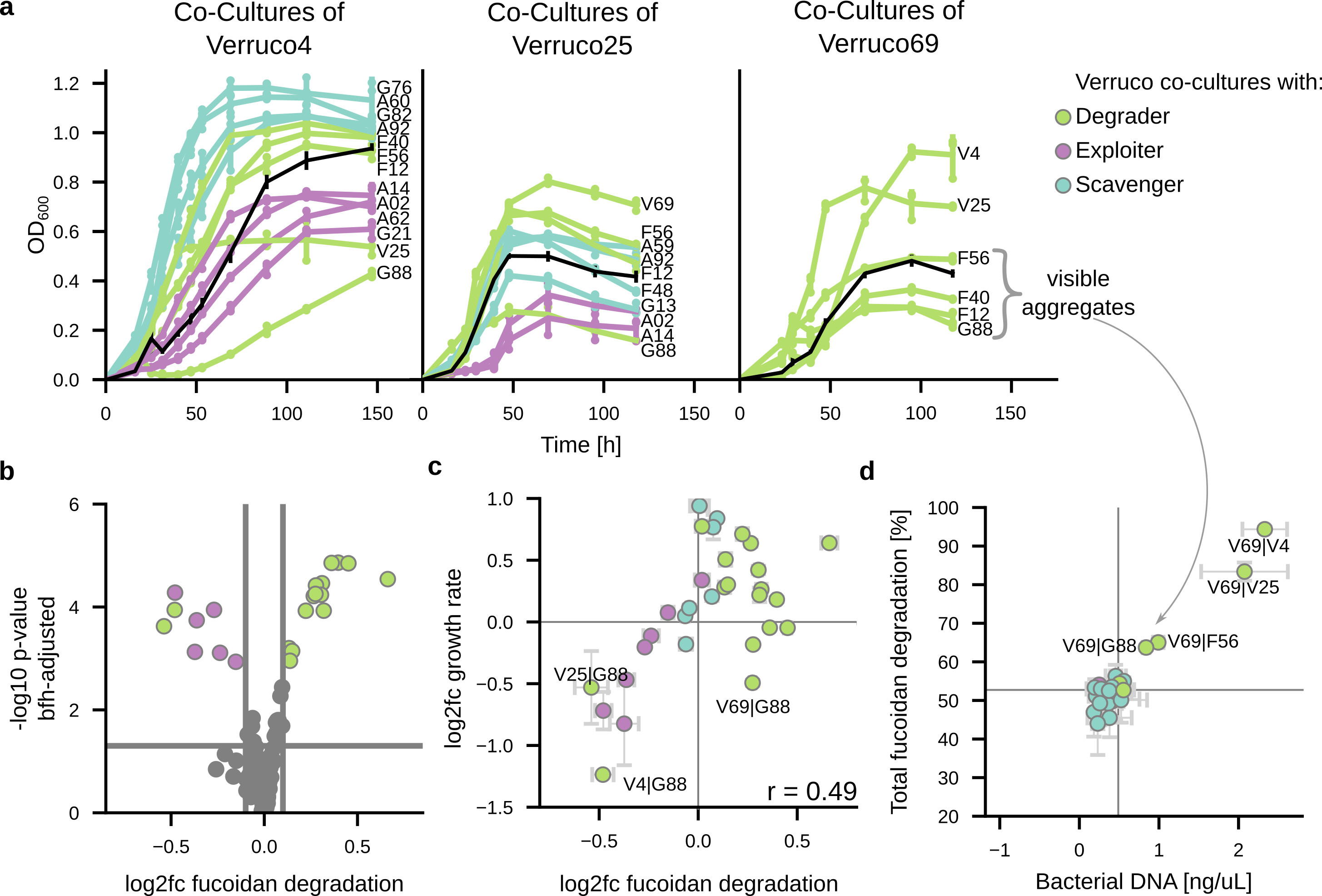

### ED7.png

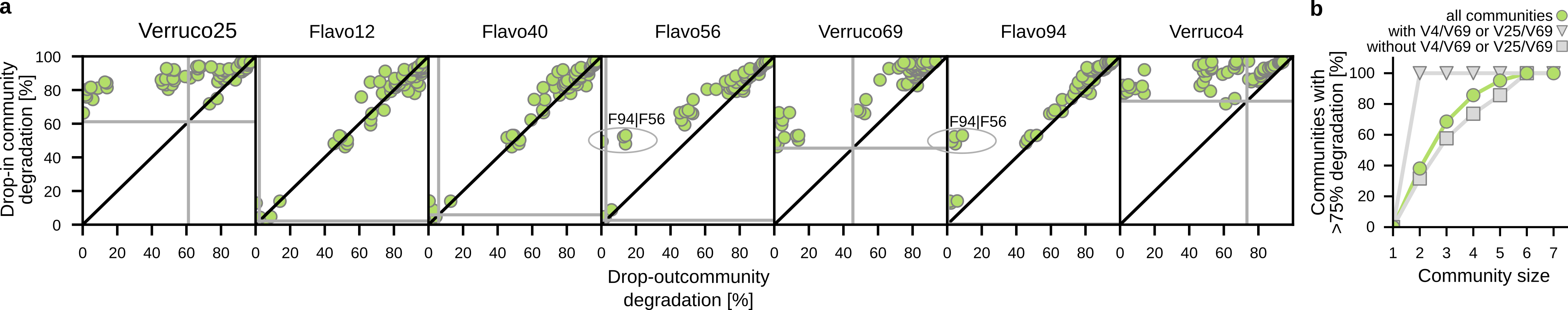

### ED7_SigmaGenes.png

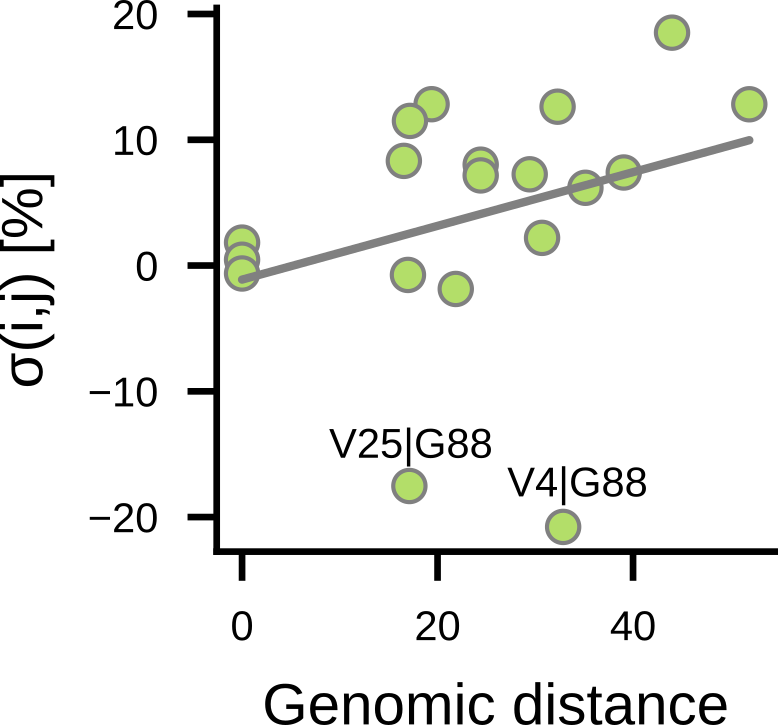

### ED9_Model.png

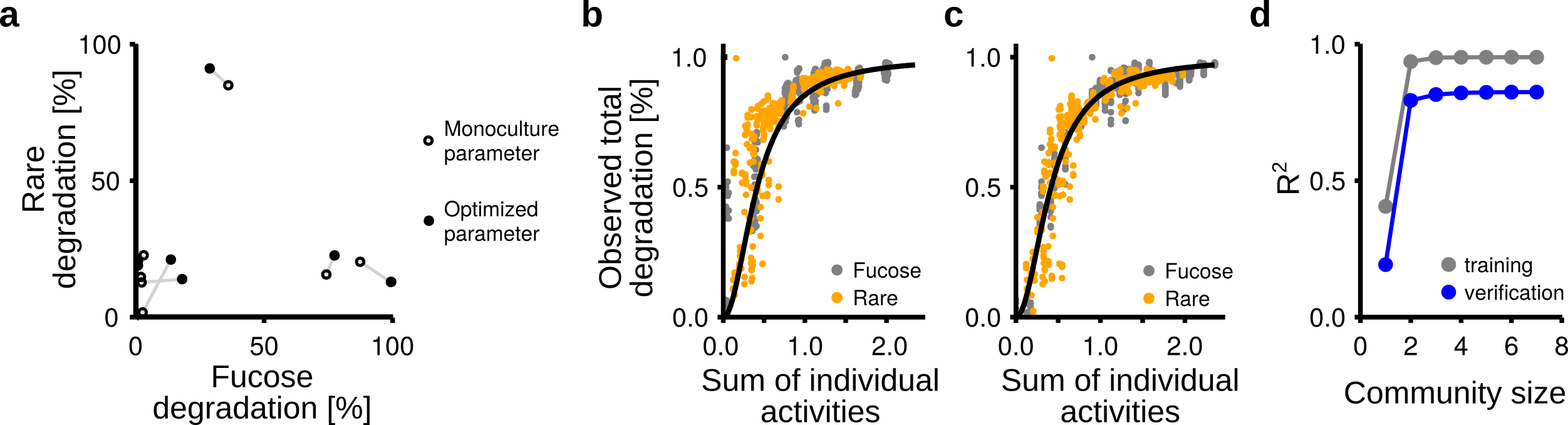

### ED10_GrowthDiversity.png

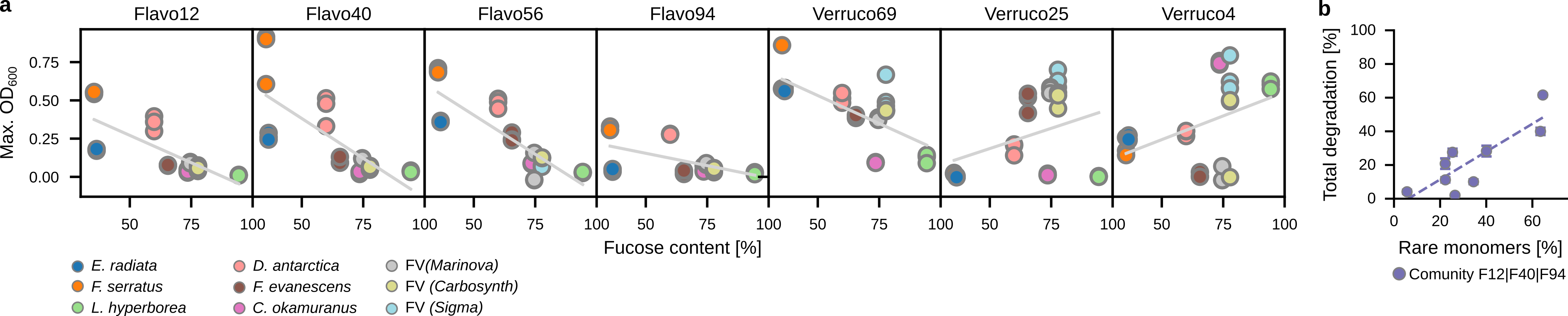
